# Oscillatory calcium release and sustained store-operated oscillatory calcium signaling prevents differentiation of human oligodendrocyte progenitor cells

**DOI:** 10.1101/2021.09.22.461371

**Authors:** Richard A. Seidman, Heba Khattab, Jessie J. Polanco, Jacqueline E. Broome, Fraser J. Sim

**Affiliations:** Neuroscience Program, Jacobs school of Medicine and Biomedical Sciences, University at Buffalo, Buffalo, NY; Department of Pharmacology and Toxicology, Jacobs school of Medicine and Biomedical Sciences, University at Buffalo, Buffalo, NY

**Keywords:** human, lentivirus, STIM1, STIM2, muscarinic receptor, metabotropic glutamate receptor

## Abstract

Endogenous remyelination in demyelinating diseases such as multiple sclerosis is contingent upon the successful differentiation of oligodendrocyte progenitor cells (OPCs). Signaling via the Gα_q_-coupled muscarinic receptor (M_1/3_R) inhibits human OPC differentiation and impairs endogenous remyelination in experimental models. We hypothesized that calcium release following Gα_q_-coupled receptor (G_q_R) activation directly regulates human OPC (hOPC) cell fate. In this study, we show that specific G_q_R agonists activating muscarinic and metabotropic glutamate receptors induce characteristic oscillatory calcium release in hOPCs and that these agonists similarly block hOPC maturation *in vitro*. Both agonists induce calcium release from endoplasmic reticulum (ER) stores and store operated calcium entry (SOCE) likely via STIM/ORAI-based channels. siRNA mediated knockdown (KD) of obligate calcium sensors STIM1 and STIM2 decreased the magnitude of muscarinic agonist induced oscillatory calcium release and attenuated SOCE in hOPCs. In addition, STIM2 expression was necessary to maintain the frequency of calcium oscillations and STIM2 KD reduced spontaneous OPC differentiation. Furthermore, STIM2 siRNA prevented the effects of muscarinic agonist treatment on OPC differentiation suggesting that SOCE is necessary for the anti-differentiative action of muscarinic receptor-dependent signaling. Finally, using a gain-of-function approach with an optogenetic STIM lentivirus, we demonstrate that independent activation of SOCE was sufficient to significantly block hOPC differentiation and this occurred in a frequency dependent manner while increasing hOPC proliferation. These findings suggest that intracellular calcium oscillations directly regulate hOPC fate and that modulation of calcium oscillation frequency may overcome inhibitory Gα_q_-coupled signaling that impairs myelin repair.

**Significance Statement:** In this study, Seidman et al. show that SOCE is a common component of ligand-based Gα_q_-coupled signaling in hOPCs and that SOCE alone is sufficient to block hOPC differentiation and drive proliferation. Therefore, SOCE blocks differentiation and pathological SOCE could contribute to myelin disease.

## INTRODUCTION

Efficient central nervous system (CNS) remyelination is contingent upon successful intercellular signaling between neurons and glia that together orchestrate the processes of oligodendrocyte progenitor cell (OPC) recruitment, differentiation, and maturation into myelin-forming oligodendrocytes (OL) ^1^. In demyelinating diseases such as multiple sclerosis (MS), the pathological environment contributes to an inhibition of OPC differentiation that limits remyelination and may contribute to subsequent permanent neurodegeneration ^2^. In support of this hypothesis, numerous quiescent OPCs are observed in chronic MS lesions that appear to be unable to undergo OL differentiation ^3,4^. Following demyelination, OPCs are exposed to a complex cellular microenvironment comprising extracellular matrix and soluble factors released by a variety of sources including infiltrating immune cells, reactive glia, and endothelial cells as well as injured axons that release both soluble growth factors and neurotransmitters. Neurotransmitters activate multiple and functionally distinct receptors expressed on OPCs and OLs that may contribute to activity-dependent or so-called adaptive myelination (reviewed in ^5^). Following demyelination, neuronal activity promotes remyelination ^6^ and this is regulated in part by AMPA and NMDA receptor signaling ^7^. As such, intracellular calcium represents a common target for these activity-dependent pathways and suggests that a diverse set of neurotransmitters can induce intracellular calcium increase in glial progenitors ^8^.

Intracellular calcium release occurs in OPCs following activation of ionotropic voltage and ligand gated channels and/or following activation of metabotropic receptors ^9^. Muscarinic acetylcholine receptors M_1_ and M_3_ are Gα_q_-coupled receptors (G_q_R) which also cause intracellular calcium release via phospholipase-Cβ (PLCβ) and inositol trisphosphate (IP_3_) second messengers ^10^. Genetic deletion of M_1/3_ receptors in OPCs delays differentiation and remyelination in animal models of experimental demyelination ^11,12^. Furthermore, pharmacological inhibition of muscarinic receptors results in accelerated OL differentiation and remyelination in the brain and spinal cord following both toxin-induced demyelination and experimental autoimmune encephalitis (EAE) ^11,13-15^. In primary human OPCs (hOPCs), M_1/3_R signaling delays differentiation *in vitro* and reduces myelin production following transplantation into hypomyelinated *shiverer* mice ^12^. OPCs have been shown to express numerous other Gα_q_-coupled receptors including endothelin (ET) subtype B (ET_B_; ENDRB) receptors ^16^, purinergic type 2Y (P2Y) receptors ^17^, histamine type 1 (H1) receptors ^18^, metabotropic glutamate receptors type 3 (mGluR_3_) and type 5 (mGluR_5_) ^19^, 5-HT serotonin receptors ^20^, sphingosine-1-phosphate receptors (S1PR) ^21^, calcium-sensing receptors (CaSR) ^22^, and adenosine 2B receptors ^23^. While the precise function of these receptors remains unclear, previous reports suggest that activation of at least some of these receptors can paradoxically promote OPC differentiation. For example, *in vitro* findings suggest that ET receptor agonists increase OPC migration and block proliferation ^16^. Stimulation of myelin basic protein (MBP) expression occurs via ET_A 24_ receptor activity, and activation of ET_B_ receptors has been found to promote remyelination in cerebellar slice culture ^25^. Likewise, conditional deletion of S1PR causes increased OPC proliferation and reduced oligodendrocyte differentiation ^26^, while CaSR null mice exhibit reduced MBP expression during development ^22^. Despite these findings, the key signaling components downstream of muscarinic receptor activation that act to delay OPC differentiation and distinguish M_1/3_R from other Gα_q_-receptors have not yet been established.

Following activation of Gα_q_-coupled receptors and calcium release from the endoplasmic reticulum (ER), calcium sensing STIM proteins in the ER membrane oligomerize and complex with ORAI channel proteins located in the plasma membrane. These calcium release activated calcium (CRAC) channels mediate store operated calcium entry (SOCE) that replenish depleted ER-stores through sarcoplasmic/endoplasmic reticulum ATPase (SERCA) pumps ^27^. Successive activation of ER-calcium release and CRAC channel influx events result in characteristic oscillatory calcium transients ^28^. Intriguingly, the frequency encoding of oscillatory calcium signaling is associated with cell-type specific activation of distinct downstream signaling and gene expression ^28,29^. In rodent OPCs, *in vitro* application of a pharmacological SOCE antagonist limits platelet derived growth factor (PDGF)-dependent proliferation ^30^. However, the mechanisms of action of these drugs are complex with additional actions on IP_3_ receptors, gap junction permeability, and transient receptor potential channels. Furthermore, while ligand-induced calcium oscillations have been observed in OL lineage cells ^19,31,32^, a specific role of calcium oscillations in human OPCs has not yet been determined.

In this study, we hypothesized that the specific oscillatory signature of ligand activated Gα_q_-receptor mediated calcium responses may be responsible for the regulation of hOPC differentiation. To this end, we identified several Gα_q_-coupled receptors expressed by hOPCs and characterized the calcium response following application of receptor subtype-specific ligands. Only activation of muscarinic acetylcholine M_1/3_ receptors or metabotropic type 5 glutamate receptors (mGluR_5_) resulted in oscillatory transients in a large fraction of OPCs and persisted for several minutes following application of the ligand. Intriguingly, these ligands also reduced hOPC differentiation. We demonstrate that SOCE downstream of M_1/3_R ligand activation is dependent on ER calcium sensors STIM1/2. More specifically, we found that both STIM proteins contribute to the magnitude of SOCE in human OPCs and that STIM2 regulates the oscillatory frequency of M_1/3_R-induced calcium responses. Knockdown of STIM2 resulted in reduced OL differentiation and prevented the anti-differentiative effects of M_1/3_R signaling in hOPCs. Lastly, using optogenetic activation of CRAC-mediated calcium flux, we demonstrate that SOCE induction is sufficient to inhibit hOPC differentiation in the absence of G_q_R signaling and that modulation of OPC proliferation and differentiation was dependent on SOCE frequency.

## MATERIALS AND METHODS

### Human CD140a/PDGFαR cell preparation and culture with Gα_q_-agonists

Fetal brain tissue samples, between 17 and 22 weeks of gestational age, were obtained from Advanced Bioscience Resources (Alameda, CA) with informed consent obtained from all donors. All research was performed in accordance with relevant guidelines/regulations. Following review by the University at Buffalo Research Subjects Institutional Review Board the tissue acquisition and research was not deemed to involve human subjects as defined under HHS regulations 45 CFR 46, 102 (f). Forebrain samples were minced and dissociated using papain and DNase as previously described ^33^. Magnetic sorting of CD140a/PDGFαR positive cells was performed as described ^34^.

### Plasmid construction, lentiviral generation, and titration

EF1α:GCaMP6s was described previously ^12^. EF1α:jRCaMP1a was cloned as follows. pGP-CMV-NES-jRCaMP1a was obtained from AddGene (#61562) ^35^. The nuclear export signal (NES) and jRCaMP1a coding region was PCR amplified: SpeI, forward−5’ AAAACTAGTGAACCGTCAGATCCGCTAG-3’;PspXI (Thermo Fisher Scientific), reverse, 5’-AAAGCTCGAGCTCTACAAATGTGGTATGGCTG-3’ (Thermo Fisher Scientific). PCR products were purified (Monarch PCR Cleanup Kit (NEB, #T1030S)) and restriction digested with unique 5’ SpeI and 3’ PspXI sites, and purified again. NES-jRCaMP1a digested PCR fragments were then ligated into lentiviral pTRIP-EF1α vector (a gift of Abdel Benraiss, University of Rochester)^36^. The plasmid pLenti-EF1α-OptoSTIM was a gift from Taeyoon Kyung (KAIST, DAEJEON, Republic of Korea) ^37^. Lentiviruses were generated as described ^38^ and titration of EF1α:GCaMP6s and EF1α:OptoSTIM was performed as previously described ^39^. Optimized multiplicity of infection (MOI) in hOPCs was determined for EF1α:GCaMP6s and EF1α:jRCaMP1a by infecting hOPCs and quantifying proportion of responsive cells following administration of 25µM Oxo-M. Optimal MOI for EF1α:OptoSTIM lentivirus was determined through analysis of optimal GFP expression in infected hOPCs.

### Calcium imaging using LV EF1a:GCaMP6s

Calcium imaging was performed as described previously ^12^. GCaMP6s infected hOPCs were seeded at 4-5 × 10^4^ cells/mL and imaged 24 hours after seeding. All experimental data were generated from independent experiments using individual naïve cultures of hOPCs from separate patient fetal samples. To avoid possible confounds relating to sequential application of small molecules, all cultures were used only once for imaging calcium responses and only a single dose was tested in each well, with two technical replicate imaging fields imaged simultaneous per well for each dose or condition tested. In dose response experiments, all doses per each drug were tested on the same experimental day for each fetal sample preparation. Bright field and fluorescence images were acquired (10x; Olympus IX51 microscope with a Exfo X-Cite or Xylis illumination and Hamamatsu ORCA-FLASHv4 camera) using µManager ^40^. All drugs were reconstituted and aliquoted within 1-2 weeks and thawed immediately prior to use. Region of interest (ROI) generation and image processing were performed using ImageJ, and analysis of calcium transients was performed in R as described ^12^.

Briefly, per cell calcium responses (raw pixel intensity) were normalized to baseline fluorescence calculated prior to addition of Gα_q_-receptor ligand (F_0_, average 5-10 frames). To define calcium peaks and characterize the properties of ligand-induced calcium oscillations, individual cell traces were loess smoothed prior to the identification of local minima and maxima in 16 second windows. Calcium peaks were defined as maxima whose amplitude increased ≥0.35 fold-change above the preceding local minima. Peak amplitude was defined as the difference in fluorescence values from the calcium peak to the preceding local minima. The number of oscillations was defined as the number of peaks identified during the recording session. A responsive cell was defined as any cell with at least one peak, and an oscillatory cell as any cell with two or more peaks. Response duration was measured from the local minima prior to the first peak oscillation to the local minima following the last peak oscillation, or to the end of the time-lapse recording if an on-going calcium peak had not yet reached baseline. For oscillatory cells, frequency was calculated by dividing the total number of peak oscillations by the response duration. Area under the curve (AUC) was calculated (*Mess* package, v0.5.6; https://cran.r-project.org/web/packages/MESS/index.html) by subtraction of F_0_, and then analyzed as integral pixel intensity above baseline beginning from the local minima preceding the first identified peak on the normalized calcium curve following ligand stimulation for the duration of the imaged response.

Raw data measurements of individual cell calcium responses were assessed for normality. Per-cell AUC and frequency measurements exhibit significant skew and were log-transformed to achieve a near normal distribution. Unless otherwise stated, a linear model was used for one-way ANOVA using drug dose as a covariate, as well as human sample to consider individual sample variability. Non-linear regression was performed using GraphPad Prism (8.0e) using a variable slope-four parameter logistic equation to model and calculate EC_50_ for each drug. For STIM KD experiments, GCaMP6s cells were transfected with STIM siRNA and imaged 24-48h post transfection.

### Assessment of store-operated calcium entry

For direct assessment of SOCE, hOPCs were infected with LV-GCaMP6s (1 MOI) and 5 × 10^4^ cells/mL were seeding in 48-well plates and imaged the following day. To remove extracellular calcium, 15 minutes prior to imaging media was replaced with HBSS without phenol red, calcium, and magnesium (Corning, #21-022) (HBSS-). For all conditions, fields were imaged with a 5 second interval for at least 15 minutes (488nm excitation, 400ms exposure). Addition of the SERCA blocker Thapsigargin (Tg) or the muscarinic agonist Oxotremorine-Methiodide (Oxo-M) with or without store operated calcium entry (SOCE) channel antagonists occurred at ∼1-2 minutes after the start of imaging. Calcium [1.2mM]) was re-introduced 10-15 minutes following addition of Tg or Oxo-M. Final concentrations of compounds were Tg [10µM] or Oxo-M [25µM], and 2-APB [50µM] or MRS-1845 [10µM] (final DMSO concentration did not exceed 0.15%, which did not influence [Ca^2+^]_i_). For quantification of SOCE, per cell calcium response curves were normalized to baseline fluorescence F_0_ (average of 5-10 frames prior to addition of Oxo-M or Tg). Normalized traces were loess smoothed to identify responsive cells exhibiting ≥0.35 fold increase in GCaMP6s fluorescence from baseline following initial Oxo-M or Tg ER-store depletion. Area under the curve (AUC) was measured for ER depletion for 10 minutes following Oxo-M or Tg ER-store depletion, and AUC for SOCE was measured for 5-8 minutes (**Fig. 1H**) or 8 minutes (**Fig. 2E**) following calcium re-addition. AUC durations were matched between conditions and were quantified. For analysis of SOCE following STIM KO, AUC of SOCE was normalized to the ER-release of control cells in matched biological replicates and quantified using repeated measures (RM) one-way ANOVA with Holm-Sidak’s post-hoc test. Maximum amplitude of ER depletion was identified as the maximal peak response >0.35 fold-change increase from baseline following Oxo-M or Tg addition and following calcium re-addition respectively. Cells which did not exhibit an initial ER-store depletion response (≥0.35 fold-change from baseline following Tg or Oxo-M addition and prior to calcium re-addition) were excluded from further analyses.

**Figure 1.**
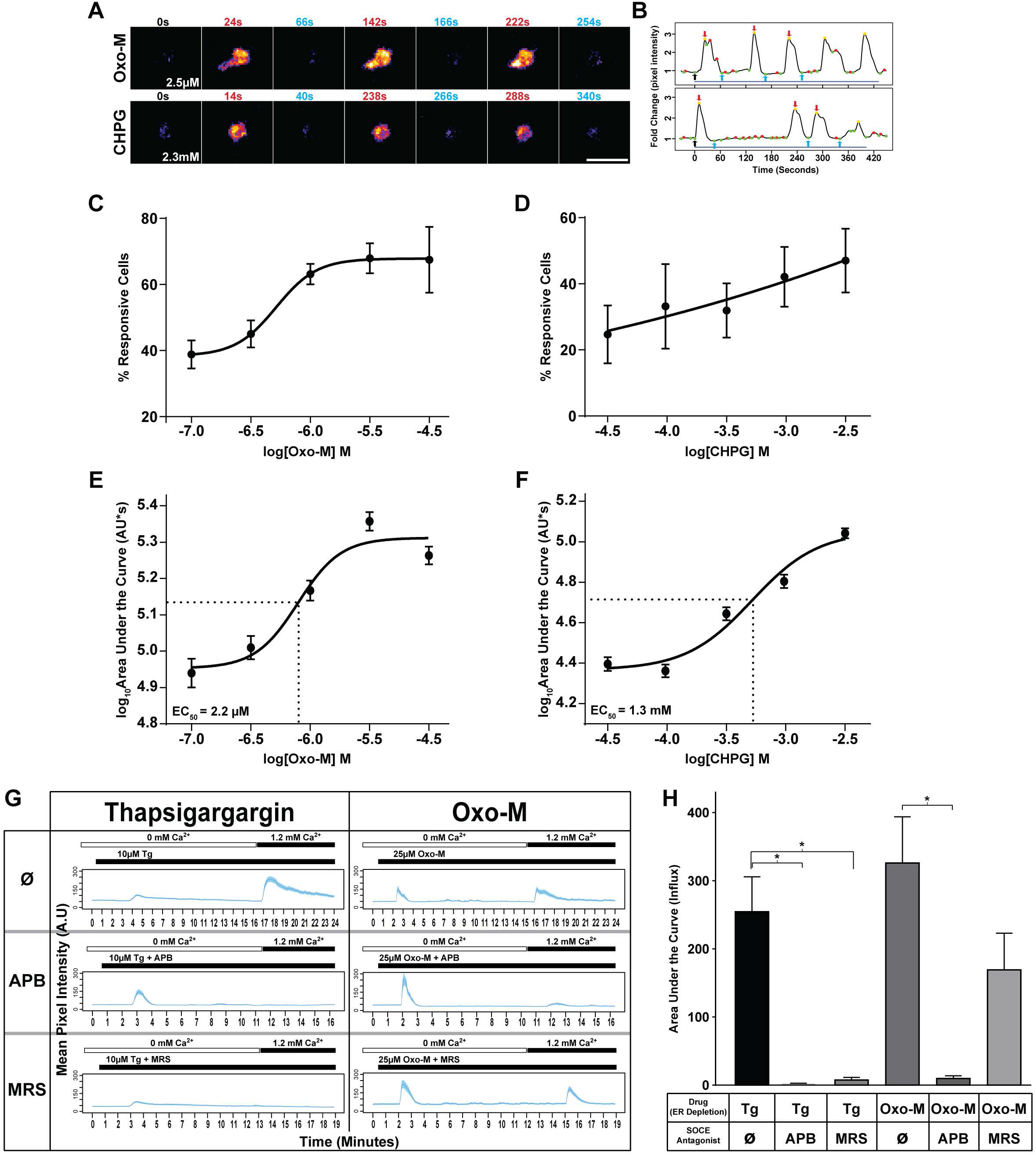
Gα_q_-receptor activation induces ER-Ca^2+^ responses and SOCE in primary human OPC. Primary fetal brain-derived PDGFαR^+^ hOPCs were infected with intracellular Ca^2+^ reporter GCaMP6s expressing lentivirus. Time-lapse microscopy of GCaMP6s fluorescence following Gα_q_-receptor agonist treatment was recorded and analyzed. **A**, pseudo-colored GCaMP6s fluorescence time-lapse images of oscillatory calcium transients following stimulation with Oxotremorine-M (Oxo-M) [2.5µM] (top) and CHPG [2.3mM] (bottom). **B**, shows corresponding normalized Ca^2+^ traces. Drug stimulation occurs at 0s (black arrow in B) and time following drug addition is indicated in each panel. Red and green dots indicated defined local maxima and minima, respectively. Gold dots indicated defined peaks. Images shown in A after drug addition correspond to calcium peaks (red) or post-peak minima (blue) and are indicated as arrows in B. Blue lines below traces indicate the calculated response duration. **C, D**, dose-response curves of the percentage of hOPCs responding to Gα_q_ ligand treatment following treatment with either oxotremorine−M (Oxo-M) or CHPG (mean ± SEM, n=3 human fetal samples, >240 cells per biological replicate). **E, F**, dose-response curve for the log_10_-transformed overall area under the curve (AUC) for [Ca^2+^]_i_ release per cell (mean ± SEM shown, n > 250 cells per dose, obtained from n=3 human fetal sample). EC_50_ for AUC was calculated by non-linear regression for each compound (variable slope, four parameters). **G**, to directly assess SOCE, GCaMP6s-expressing hOPCs were cultured in calcium-free conditions for 5-10 minutes prior to time-lapse microscopy. Following a 1-2m baseline, cells were treated with thapsigargin (Tg) or Oxo-M, followed by re-addition of calcium containing solution [1.2mM] to stimulate SOCE (green dashed lines) in the presence (middle and bottom traces) or absence (top trace) of SOCE antagonists. Calcium traces showing mean pixel intensity ± SEM (blue shading) of ≥ 44 cells from a single biological replicate. Timings for addition of Tg, Oxo-M and Ca^2+^-containing solution are indicated by horizontal bars above each plot. **H**, quantification of SOCE was performed by analysis of AUC of calcium influx following calcium re-addition (mean ± SEM; Tg: n=2 human fetal sample preparations; OxoM: n=3 human fetal sample preparations). * p<0.05, RM one-way ANOVA with Holm-Sidak’s post-hoc test. Scale: 25µm.

**Figure 2.**
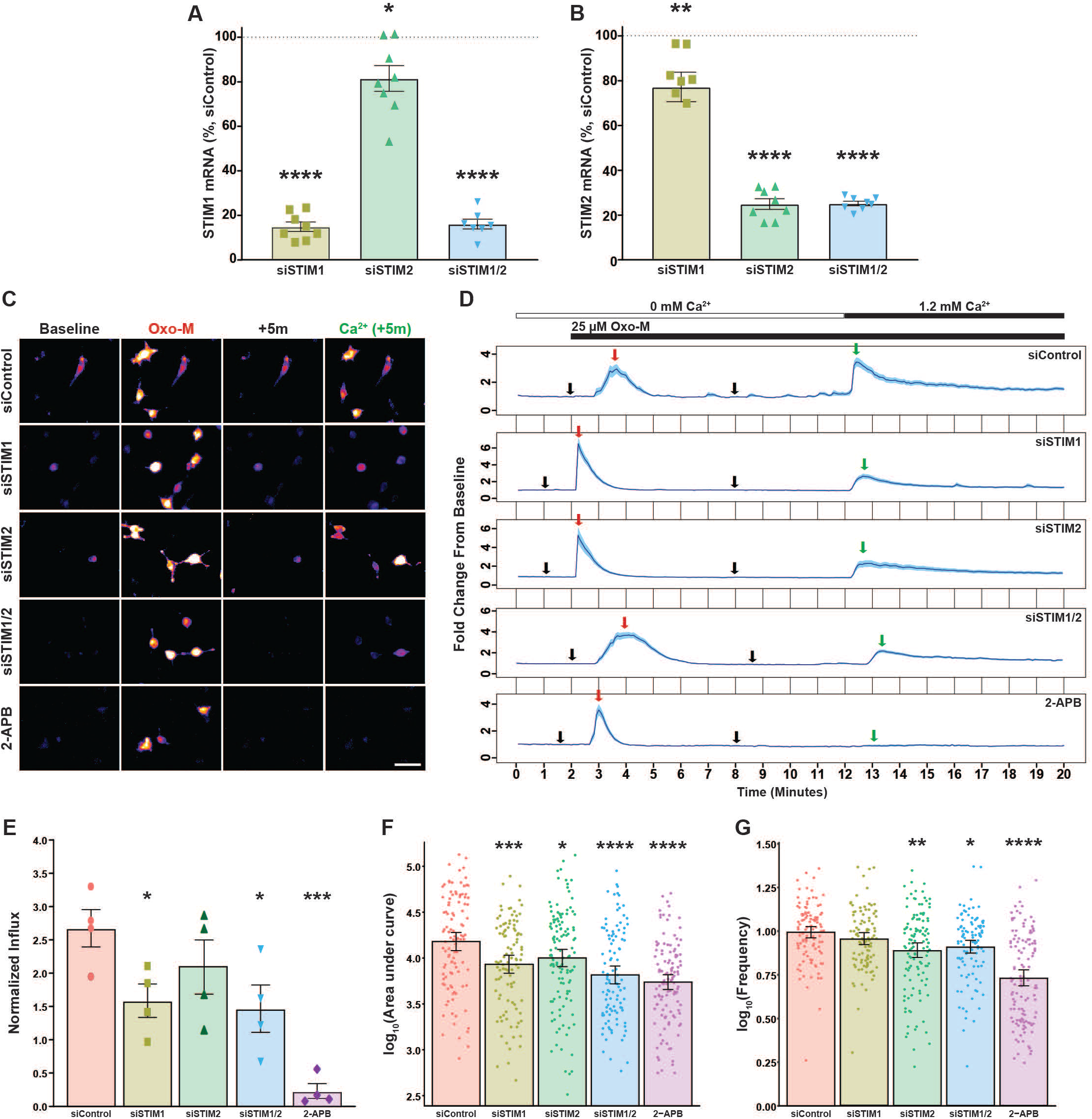
STIM1/2 calcium sensors are required for muscarinic-induced SOCE and [Ca^2+^]_i_ response in human OPCs. hOPCs were transfected with STIM1 and/or STIM2, or scrambled siRNA, cultured in mitogen-containing media, and RNA was extracted after 24-48 hours. For calcium imaging experiments, hOPCs were initially infected with GCaMP6s lentivirus and then transfected with STIM RNAi or scrambled control siRNA prior to time-lapse microscopy. RT-qPCR analysis of STIM1 (**A)** or STIM2 (**B**) mRNA expression relative to scrambled control transfected hOPCs following STIM1, STIM2 or STIM1/2 RNAi knockdown (STIM KD) (mean ± SEM, n=7-8 human samples). Control normalized mean ± SEM, one-sample t-test, *p<0.05, **p<0.01, ****p<0.0001. **C**, representative pseudo-colored fluorescence time-lapse images of peak Oxo-M-induced calcium responses in siRNA transfected hOPCs cultured in calcium-free conditions. Images represent peak Oxo-M [25µM] induced ER-store depletion (red text and arrows in **C** and **D** respectively) and SOCE following calcium re-addition (green text and arrows in **C** and **D** respectively). **D**, calcium traces of Oxo-M induced calcium response following siRNA transfection showing normalized mean pixel intensity ± SEM (blue shading) of n ≥ 30 cells per condition from one representative biological replicate. Timings for addition of Oxo-M and Ca^2+^-containing solution are indicated by horizontal bars above plots. **E**, SOCE was measured as the total area under the curve following calcium re-addition normalized to ER-release of control cells in the matched biological replicate. STIM1/2 KD and pharmacological SOCE inhibition both resulted in a significant reduction in muscarinic-induced SOCE relative to control hOPCs. Cells were analyzed across 1-2 imaging fields in a single well per each condition and all responses were averaged per condition for each biological replicate. Data are presented as mean ± SEM of averaged cell responses per each of four independent experiments, with >140 total cells quantified per each condition, n=4 independent human fetal sample preparations. RM one-way ANOVA with Holm-Sidak’s post-hoc test, * p<0.05, ** p<0.01). **F−G**, the net effect of pharmacological SOCE inhibition or STIM KD on Oxo-M-induced increased [Ca^2+^]_i_ was determined in calcium-containing normal growth media by time-lapse microscopy. STIM1 KD or STIM2 KD hOPCs displayed a reduction in Oxo-M-induced calcium responses as denoted by reduced average AUC (**F**) as well as a reduction in oscillatory frequency relative to control hOPCs (**G**). Cells were analyzed across two imaging fields in a single well per each condition and all responses were averaged per condition for each biological replicate. Data represent log_10_-transformed mean ± SEM of per cell responses from four independent experiments (n=4 independent human fetal sample preparations), with >100 total cells quantified per each condition. * p<0.05, ** p<0.01, *** p <0.001, ****p<0.0001, linear model with Tukey’s HSD posttest. Scale: 50 µm.

### siRNA transfection

hOPCs were seeded at 5 × 10^4^ cells/mL and Lipofectamine RNAi max (Invitrogen, #13778-150) was used to transfect cells according to the manufacturer’s protocol with minor alterations. Briefly, stealth RNAi negative control duplex medium GC content (Invitrogen, #12935300) or three combined stealth RNAis against STIM1 (Invitrogen, ID#’s: HSS110308, HSS110309, HSS186144) and/or STIM2 (Invitrogen, ID#’s: HSS183972, HSS183973, HSS183974) were pooled and used for transfection. Stealth RNAi (100 nM total RNAi per well) was used to transfect hOPCs per each condition with 1.5% v/v Lipofectamine. For calcium imaging, cells were plated in 48-well plates and imaged 24 hours post-transfection. In differentiation experiments, mitogen containing media was removed 24 hours post-transfection (experimental day 0) and replaced with media without mitogens. Oxo-M [25µM] was added on experimental day 0 upon removal of growth factors and on experimental day 2 during a half media change. hOPCs were fixed on experimental day 3-4 following live staining with mouse × O4 Hybridoma primary antibody [1:25]. Immuno-cytochemical analysis was performed as described previously ^33^. For analysis of STIM expression, cells were plated at 5 × 10^4^ cells/mL 6-well plates and transfected after 24 hours. RNA was extracted 24-48 hours post-transfection for cDNA synthesis and quantitative RT-PCR analysis.

### Quantitative RT-PCR analyses

Total RNA was extracted (Omega Biotek) and cDNA was synthesized using SuperScript III reverse transcriptase (Invitrogen, Carlsbad, CA). Human-specific primers for SYBR green-based PCR were designed (**Table 1**). Samples were run in triplicate and gene expression calculated by ΔΔC_t_ method using GAPDH as a reference. Statistical significance was tested on log_2_-transformed data normalized to control using a one-sample t-test assuming equal SD and compared to a hypothetical value of 100 (GraphPad Prism 8.0e).

**Table 1:**
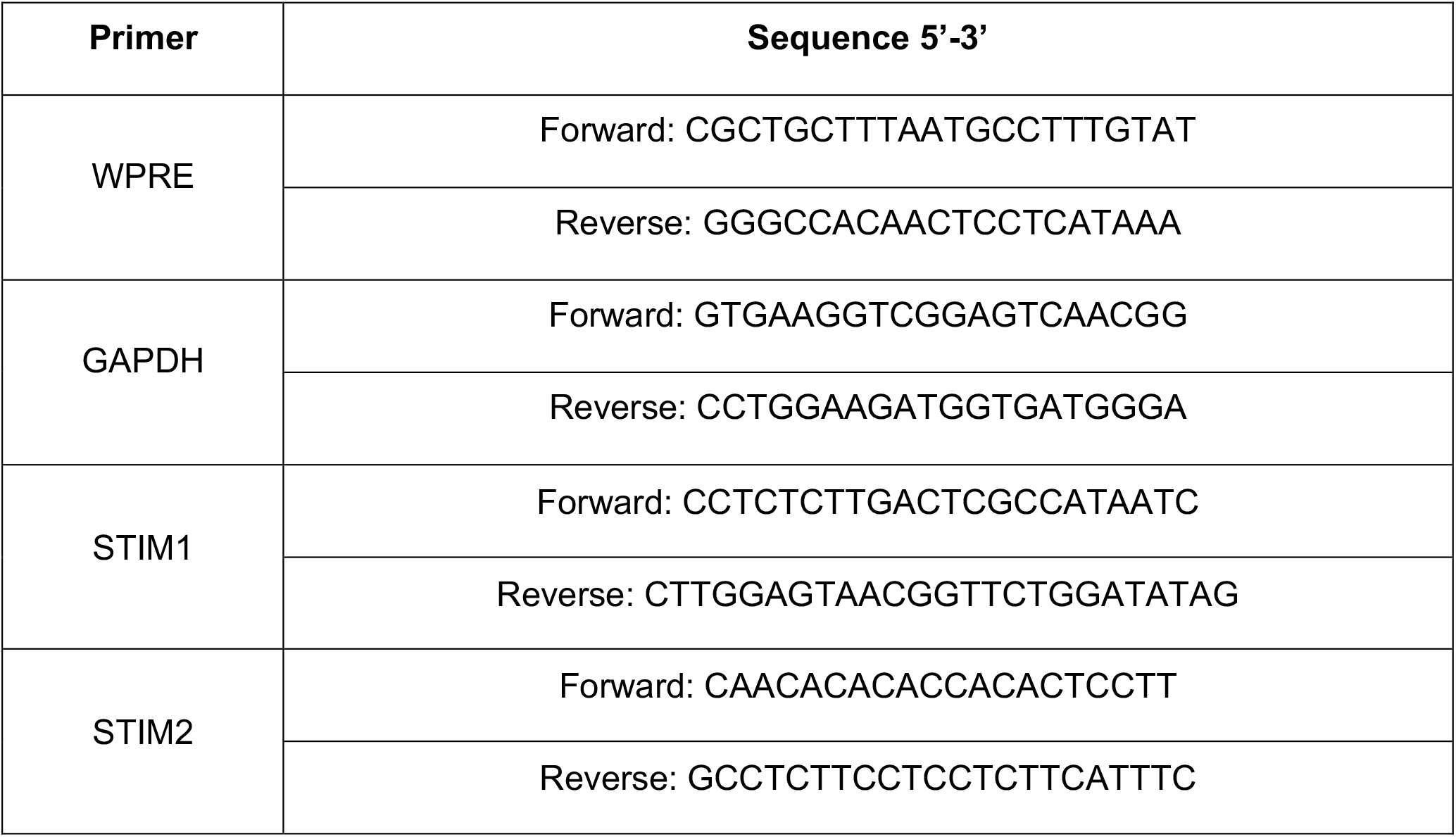
qPCR primers.

### Assessment of long-term OptoSTIM-induced SOCE

For all long-term experiments, hOPCs were seeded at a density of 2.5-3.0 × 10^5^ cells/mL into poly-ornithine and laminin coated ibidi tissue culture treated µ-slide VI 0.4 (iBIDI, #80606) in mitogen containing Livelight ND media to reduce phototoxicity ^41^. Cells were then imaged in an on-stage ibidi environmentally controlled chamber to maintain 37ºC, 5.0% CO2, and >60% humidity for the duration of each imaging experiment. In long-term calcium imaging experiments, hOPCs were infected with jRCaMP1a (1 MOI) and OptoSTIM (1 MOI) lentiviruses. Two fields per each of 3 µ-slide channels were then imaged at 10x with 594 nm excitation every 20 seconds for 3 hours to measure jRCaMP1a fluorescence calcium responses. After a 30-minute no-stimulation period and for the remainder of imaging, hOPCs were stimulated with 488 nm excitation (every 5 or 30 minutes with 330 and 2000 ms illumination times respectively) to induce SOCE calcium responses. A custom beanshell script was designed and implemented to facilitate multi-channel and multi-position imaging and provide an interface between the motorized stage, fluorescent shutter (Prior), and µManager software. The complete imaging script is available upon request. Analysis of OptoSTIM-induced calcium responses was conducted by implementing the automated tracking plugin TrackMate in ImageJ ^42^. Mean fluorescence intensity data within the region of interest for all cells tracked across all frames was exported and subsequently analyzed in R. Briefly, a baseline curve was generated by fitting minima points under each calcium trace using the “spc-Rubberband” function (*hyperSpec* package). Traces were first normalized by division of the baseline curve and loess fitted to identify responsive cells with ≥12.5% increase in jRCaMP1a fluorescence within 2 minutes following the first OptoSTIM 488nm pulse. For quantification of AUC, each tracked cell’s mean jRCaMP1a fluorescence values per each time-point were normalized by subtraction of the baseline curve and AUC was calculated for the duration of the stimulation period. Calcium traces are presented as fold change in jRCaMP1a fluorescence from baseline, averaged across all cells per each time-point with SEM calculated. In differentiation and proliferation experiments, OptoSTIM infected hOPCs (1 MOI) were seeded into µ-slides and bright field imaged for 2−3 days. Following a 10-12 hour baseline period, OptoSTIM activation was initiated with 488nm stimulation at different frequencies (every 5 or 30m with 330 or 2000ms illumination times respectively) for the remainder of imaging. After imaging, cells were immunostained for O4 and the number of O4^+^ and GFP^+^ cells were counted and analyzed. Repeated measures (RM) one-way ANOVA was employed with Holm-Sidak’s post-hoc test for quantification of the proportion of O4^+^ to GFP^+^ cells and quantification of morphological branching. In proliferation experiments, cell density was counted at the onset of blue light stimulation and mitotic events were identified and counted throughout the stimulation period. RM one-way ANOVA with Holm-Sidak’s post-hoc test was implemented to quantify significance in proliferative events normalized to initial densities for each condition.

## RESULTS

### Gα_q_-receptor activation induces ER-Ca^2+^ responses and SOCE in hOPC

We hypothesized that Gα_q_-coupled receptors would exert distinct effects on intracellular calcium responses in hOPCs and that these responses would correlate with their functional effects. Using RNA-seq of human primary PDGFRα-isolated OPCs, we determined that hOPCs expressed a diverse set of 122 GPCRs including 14 GPCRs that couple via Gα_q_ proteins (GPCRs “GO:0004930” and PLC activating Gα_q_-coupled “GO:0007200”, respectively). To determine the patterns of Gα_q_-mediated calcium responses in hOPCs, we used lentivirus to express the intracellular calcium reporter GCaMP6s hOPCs. We next exposed independent cultures to receptor selective ligands at 10-fold the reported EC_50_ concentration and measured resultant calcium responses (**Fig. 1** and **Table 2**). All ligands induced a detectable increase in intracellular calcium concentration measured by GCaMP6s fluorescence. However, consistent with receptor heterogeneity, the majority of ligands tested increased calcium in only a small fraction of GCaMP6-expressing hOPCs (between 1-30%). In contrast, activation of M_1/3_R and mGluR_5_ receptors by Oxo-M and CHPG, respectively, elicited robust responses in >46% of hOPCs (n = 3 human samples). Both agonists induced a dose-dependent increase in the proportion of responding cells (**Fig. 1C-D**). The extent of total calcium flux was determined by measuring the area-under-curve (AUC). AUC was defined as the area of the response above baseline beginning at the initial response until the end of the imaging session (see methods). Raw AUC values were log_10-_normalized to achieve normality and displayed as mean ± SEM of pooled responses from >250 cells quantified per each dose from three independent experiments using three independent human samples. Both agonists induced a clear dose-dependent intracellular calcium response with EC_50_ determined as 2.2 µM and 1.3 mM for Oxo-M and CHPG respectively (**Fig. 1E-F**).

**Table 2:** Human OPC-expressed Gα_q_-coupled receptors differentially regulated oscillatory intracellular calcium fluctuations. RNA-seq data from human and mouse ^46^ was used to identify Gα_q_-coupled receptors expressed by OPCs. FPKM values shown (mean ± SEM, n = 2 independent human fetal sample preparations). Ligands activating each receptor were identified to determine the calcium response characteristics in hOPCs. hOPCs were infected with GCaMP6s-expressing virus and exposed to various Gα_q_-coupled receptor ligands at doses approximately 10-fold larger than reported EC_50_ values (or previously effective doses in rodent OPCs*). Calcium responses were recorded for at least 10 minutes and analyzed as described. The percentage of responding cells (at least 1 peak response ≥ 0.35 fold increase from baseline on normalized calcium trace following ligand addition), maximum response amplitude (fold change from baseline), duration of response (minutes) and percentage of responding cells displaying an oscillatory calcium response (cells with ≥ 2 identified peak responses) were determined. While Gα_q_-coupled LPA receptors LPAR1 and LPAR4 were detected in hOPCs, due to the lack of specific ligands we did not test these receptors. Mean ± SEM of total % responsive cells from n=2-3 independent human fetal sample culture preparations (Oxo-M, CHPG and ET-1 data: n=3 independent fetal sample culture preparations; data for all additional ligands: n=2 individual fetal sample culture preparations). Abbreviations: **2-FLY**, 2-furoy-Leu-Ile-Gly-Arg-Leu-Orn-NH2; **2-PED**, 2-pyridylethylamide dihydrochloride; **5-HT**, 5-hydroxytryptamine; **Bay**, Bay 60-6583 2-[[6-Amino-3 5-dicyano-4-[4-cyclopropylmethoxy phenyl]-2-pyridinyl]thio]-acetamide; **CHPG**, 2-Chloro-5-hydroxyphenylglycine sodium salt; **ET-1**, Endothelin-1; **MRS 2365**, [[(1R 2R 3S 4R 5S)-4-[6-Amino-2-(methylthio)-9H-purin-9-yl]-2 3-dihydroxybicyclo[3.1.0]hex-1yl]methyl] diphosphoric acid mono ester trisodium salt; **NAGly**, N-Arachidonoylglycine; **NF 546**, 4 4’-(Carbonylbis(imino-3 1-phenylene-carbonylimino-3 1-(4-methyl-phenylene)carbonylimino))-bis(1 3-xylene-alpha alpha’-diphosphonic acid; **Oxo-M**, Oxotremorine methiodide; **PF9**, 2-(Phenylethynyl)adenosine-5’-triphosphate tetrasodium salt; **TC-G**, 4-Methoxy-N-[[[2-trifluoromethyl)[1 1’-biphenyl]-4-yl]amino]carbonyl]-3-pyridinecarboxamide; **TRAP-6**, Thrombin receptor activating peptide 6.

| Gene Symbol | FPKM (hOPC;<br>mOPCs) | Agonist/<br>Doses<br>tested | Reported<br>EC<sub>50</sub> | % Responsive | Maximum<br>amplitude | Duration<br>(min) | %<br>Oscillatory |
| --- | --- | --- | --- | --- | --- | --- | --- |
| CHRM3 | 16.3 ± 2.0;<br>0.27 ± 0.11 | Oxo-M<br>[25 µM] | 2.5 µM<sub>75</sub> | 67.6 ± 10.0 | 2.17 ± 0.06 | 6.1 ± 0.8 | 58.3 ± 20.5 |
| CHRM1 | 4.0 ± 0.9;<br>2.06 ± 0.66 |  |  |  |  |  |  |
| GRM5 | 1.2 ± 0.004;<br>13.23 ± 3.63 | CHPG<br>[7.5 mM] | 750µM<sub>76</sub> | 46.9 ± 9.7 | 0.87 ± 0.03 | 6.7 ± 0.1 | 48.6 ± 4.1 |
| S1PR1 | 3.1 ± 0.3;<br>34.32 ± 8.80 | TC-G 1006<br>[350 nM] | 35 nM<sub>77</sub> | 27.8 ± 2.3 | 0.6 ± 0.02 | 2.3 ± 0.2 | 24.8 ± 14.1 |
| F2R | 26.7 ± 0.1;<br>23.54 ± 3.31 | TRAP-6<br>[8.0 µM] | 800 nM<sub>78</sub> | 21.2 ± 13.3 | 1.0 ± 0.10 | 2.8 ± 0.5 | 38.9 ± 7.3 |
| GPR17 | 3.6 ± 1.4;<br>440.98 ± 42.13 | PF9<br>[400 pM] | 36 pM<sub>79</sub> | 19.3 ± 0.7 | 0.6 ± 0.01 | 2.3 ± 0.3 | 17 ± 10.3 |
| EDNRB | 17.3 ± 01;<br>292.72 ± 10.54 | ET-1<br>[150 nM] | 10 nM<sub>80</sub> | 16.8 ± 4.3 | 1.03 ± 0.09 | 2.4 ± 0.3 | 28.8 ± 5.6 |
| P2RY1 | 1.0 ± 0.2;<br>4.94 ± 1.46 | MRS 2365<br>[12 nM] | 1.2 nM<sub>81</sub> | 14.6 ± 8.4 | 2.9 ± 0.20 | 9.3 ± 0.9 | 48.3 ± 0.3 |
| HTR2B | 0.8 ± 0.2;<br>0.12 ± 0.02 | 5-HT<br>[100 µM] | * 10 µM<sub>20</sub> | 13.0 ± 12.4 | 0.7 ± 0.08 | 2.1 ± 0.1 | 14.3 ± 14.3 |
| F2RL1 | 1.3 ± 0.02;<br>0.10 ± 0.10 | 2-FLY<br>[1.0 µM] | 100 nM<sub>82</sub> | 8.5 ± 1.3 | 1.6 ± 0.16 | 11.2 ± 0.2 | 63.7 ± 7.0 |
| HRH1 | 0.4 ± 0.1;<br>5.01 ± 0.49 | 2-PED<br>[560 µM] | 56 µM<sub>83</sub> | 7.4 ± 3.7 | 0.9 ± 0.06 | 3.1 ± 0.1 | 29.2 ± 4.2 |
| GPR18 | 0.9 ± 0.4;<br>0.10 ± 0.10 | NAGly<br>[10 µM] | 1 µM<sub>84</sub> | 5.0 ± 3.9 | 1.3 ± 0.23 | 6.3 ± 3.5 | 21.8 ± 21.8 |
| ADORA2B | 0.6 ± 0.2;<br>8.79 ± 0.88 | Bay<br>[5.8 µM] | 580 nM<sub>85</sub> | 2.7 ± 2.7 | 0.4 ± 0.40 | 1.4 ± 1.4 | 12.5 ± 12.5 |
| P2RY11 | 0.3 ± 0.1;<br>— | NF 546<br>[5.0 µM] | 500 nM<sub>86</sub> | 1.1 ± 1.1 | 0.28 ± 0.28 | 0.9 ± 0.9 | 0.0 ± 0.0 |

The calcium responses to Gα_q_-specific agonists were further classified into monotonic and oscillatory responses (**Table 2**). Intriguingly, of those ligands which resulted in calcium transients in <30% of cells, we often observed that the majority of responses were monotonic and typically short-lived returning to baseline within 3 mins following agonist application. In contrast, agonists for M_1/3_R and mGluR_5_ receptors resulted in prolonged oscillatory calcium transients which persisted for over nine minutes (9.5 ± 2.0 and 11.0 ± 0.6 mins, for Oxo-M and CHPG respectively; mean ± SEM, n=3 human samples). For all doses tested, Oxo-M elicited oscillatory calcium signals in ∼50% of all responsive cells (**Supplementary Fig. S1A**). In comparison, cells responsive to CHPG treatment exhibited a significant increase in the percentage of oscillatory responsive cells at higher concentrations (75µM vs 2300µM, RM one way ANOVA with Holm-Sidak’s post-hoc test, p<0.05) (**Supplementary Fig. S1B**). The number of oscillations did not differ significantly between Oxo-M nor CHPG concentrations tested (**Supplementary Fig. S1C, D**). There was a significant increase in maximum amplitude with Oxo-M dose (main effect, p=0.03), which was not observed in CHPG treated cells (**Supplementary Fig. S1E, F**). Both ligands induced oscillatory calcium signals with an average frequency of ∼10 mHz (**Supplementary Fig. S1G, H**). As such, the calcium responses following activation of Gα_q_-coupled receptors were distinguishable from one another, and the similarity of prolonged oscillations in the calcium response following mGluR_5_ agonist treatment to M_1/3_R suggested that a common downstream signaling pathway may be engaged compared to other ligands.

As M_1/3_R activation leads to a profound blockade of hOPC differentiation ^12^, we hypothesized that the prolonged oscillatory calcium response may underlie this effect and that mGluR_5_ activation would therefore lead to a similar inhibition of differentiation. In addition, we predicted that EDNRB activation, which instead elicits a minimal and transient calcium response, would not influence hOPC differentiation. To test this hypothesis, we treated hOPCs with ET-1 or CHPG in the absence of mitogens and determined the effect on O4^+^ oligodendrocyte (OL) differentiation after 3-4 days (**Supplementary Fig. S2**). 150 nM and 1.5 µM ET-1 did not influence the proportion of O4^+^ OLs (RM One-way ANOVA with Holm-Sidak’s post-hoc test, p=0.64, n = 2 human samples) (**Supplemental Fig. S2A, B**). Furthermore, ET-1 did not alter the generation of complex branched O4^+^ oligodendrocytes (p=0.59) (**Supplemental Fig. S2C**). In contrast, treatment with the mGluR_5_ agonist CHPG [1.3mM] significantly attenuated O4^+^ OL differentiation from 42 ± 4% to 32 ± 5% (**Supplemental Fig. S2D, E**); RM One-way ANOVA with Holm-Sidak’s post-hoc test, p=0.01, n=5 human samples). Furthermore, CHPG treatment significantly reduced complex branch formation in differentiated O4^+^ immature oligodendrocytes from 27 ± 4% to 18 ± 5% (p=0.006) suggesting blockade of OL maturation (**Supplemental Fig. S2F**). These data suggest that distinct patterns of calcium signaling occur following Gα_q_-receptor activation and that calcium signaling downstream of Gα_q_-receptor activation may act to block hOPC differentiation and maturation.

### SOCE is activated following M_1/3_R agonist stimulation

Oscillatory calcium responses can be sustained following endoplasmic reticulum (ER) store release by several mechanisms including calcium recycling and, importantly, by store-operated calcium entry (SOCE) through the activation of calcium release activated channels (CRAC) (for review see^27^). We hypothesized that CRAC channels were activated following M_1/3_R Gα_q_-receptor activation and store depletion in hOPCs. Calcium responses in hOPCs were measured following infection with lentivirus encoding calcium reporter GCaMP6s ^12^. To investigate whether SOCE occurs following ER Ca^2+^ depletion in hOPCs, we treated cells in the absence of extracellular Ca^2+^ with thapsigargin (Tg) [10µM], a sarco-endoplasmic reticulum Ca^2+^-ATPases inhibitor (SERCA) blocker ^43^ (**Fig. 1G & Supplemental Fig. S3A**). After the initial peak in calcium due to store depletion, we observed a clear SOCE associated peak following the reintroduction of calcium into the media. Pre-incubation with CRAC channel inhibitors 2-APB ^44^ or MRS-1845 ^45^ substantially reduced SOCE in hOPCs (**Fig. 1G & Supplemental Fig. S3C, E**). We quantified SOCE as the area under the calcium curve (AUC) following calcium re-addition (n = 2 independent human sample preparations with ≥ 44 cells quantified per condition/replicate). We observed substantial SOCE following ER depletion with Tg (254.5 ± 51.3 AUC arbitrary units) which was nearly abolished by pre-incubation with 2-APB or MRS-1845 (**Fig. 1H;** RM one-way ANOVA with Holm-Sidak’s post-hoc test, p<0.05; 2-APB: 0.2 ± 0.02; MRS: 7.1± 4.3). There were no differences in the initial cytosolic calcium release following Tg induced ER depletion in each condition (RM one-way ANOVA, AUC, p=0.42). Together these results were consistent with robust SOCE in hOPCs following ER-depletion that could be blocked by pharmacological CRAC channel antagonists.

To test the hypothesis that M_1/3_R activation leads directly to SOCE, we assessed SOCE following muscarinic agonist treatment. Similar to Tg treatment, Oxo-M [25µM] induced robust SOCE following calcium reintroduction (**Fig. 1G-H & Supplemental Fig. S3B;** 326 ± 67.72 AUC, n = 3 human sample, with ≥48 cells quantified per condition/replicate). Likewise, pre-incubation with 2-APB similarly resulted in a substantial reduction of SOCE (**Fig. 1G-H & Supplemental Fig. S3D;** RM one-way ANOVA with Holm-Sidak’s post-hoc test, p=0.037; Oxo-M+2-APB: 9.8 ± 3.9). While not significant, we also observed a ∼50% reduction in SOCE following pre-incubation with MRS (**Fig. 1G-H & Supplemental Fig. S3F**, Oxo-M+MRS: 169.3 ± 53.8, p=0.1). As above, there were no differences in the magnitude of intracellular calcium release following ER-depletion between all Oxo-M conditions tested (RM one-way ANOVA, p=0.2). Together, these results show that robust SOCE occurs following M_1/3_R induced signaling in hOPCs.

### STIM1/2 expression is necessary for muscarinic receptor induced SOCE

The permeability of CRAC channels are regulated by STIM calcium sensor proteins ^27^. RNA-seq analyses of human OPCs revealed expression of both calcium sensors STIM1 and STIM2 (STIM1: 8.69 ± 0.004 and STIM2: 30.0 ± 0.56; FPKM ± SEM, n=2) and was similar to that observed previously in mouse OPCs ^46^ (Stim1: 13.9 ± 1.4 and Stim2: 14.9 ± 0.7). We hypothesized that the calcium sensors STIM1 and STIM2 would be directly involved in regulating calcium influx through CRAC channels following Gα_q_-receptor activation in hOPCs. To test this hypothesis, we used siRNA targeting STIM1 and/or STIM2 coupled with calcium imaging. We observed a 70-85% decrease in mRNA expression of STIM1 (**Fig. 2A;** one sample t-test p<0.0001) and STIM2 (**Fig. 2B;** STIM2 & STIM1/2: p<0.0001) following STIM knockdown (KD) in hOPCs. Interestingly, we also observed a small but significant decrease in STIM1 expression (p<0.05) following STIM2 siRNA transfection, as well as a reduction in STIM2 expression following STIM1 siRNA (p<0.01). To directly assess the role of STIM proteins in muscarinic receptor-induced SOCE, we assessed calcium influx following extracellular calcium removal and re-addition in the context of STIM1/2 KD and Oxo-M treatment. As shown above we identified robust SOCE downstream Oxo-M stimulation which was effectively blocked by CRAC antagonist 2-APB (n=4 independent human fetal sample preparations, RM one-way ANOVA with Holm-Sidak’s post-hoc test, p=0.003). STIM1 and STIM1/2 KD significantly reduced SOCE by ∼40% (STIM1: p=0.047; STIM1/2: p = 0.041) while STIM2 had no effect (p=0.19) (**Fig. 2C-E)**. These results suggest that STIM1 contributes to muscarinic-induced SOCE and is a key component of the endogenous hOPC CRAC apparatus. As STIM1/2 silencing may influence endogenous ER-store calcium content, we analyzed the total calcium flux (AUC) and maximum amplitude of ER calcium store depletion in response to Oxo-M following STIM1/2 KD in the absence of extracellular calcium. RM one-way ANOVA revealed no significant differences in either the AUC (**Supplemental Fig. S4A**, Holm-Sidak’s post-hoc tests, p=0.1) or maximum amplitude (**Supplemental Fig. S4B**, p>0.4) of ER store release following STIM KD compared to control cells, indicating that STIM silencing did not influence basal ER calcium store content or the extent of ER-calcium depletion following G_q_ stimulation.

We next sought to investigate the effects of STIM1 and STIM2 KD and their roles in regulation of ligand induced Ca^2+^ oscillations in the presence of extracellular calcium. While STIM1 KD induced a greater reduction in Ca^2+^ release following Oxo-M treatment, knockdown of STIM1 or STIM2 significantly attenuated the magnitude of Ca^2+^ release (**Fig. 2F**; linear model with Tukey’s HSD posttest, p<0.0001, n=4 human samples). Intriguingly, STIM2 KD also resulted in a significant reduction in the frequency of Ca^2+^ oscillations (**Fig. 2G**; STIM2: p= 0.002; STIM1/2: p=0.0323) while STIM1 KD had no effect (STIM1: p = 0.7236). We also observed that neither STIM1 KD nor STIM2 KD altered the percentage of oscillatory responsive cells (**Supplemental Fig. S5A**, RM one-way ANOVA, p=0.45) or the maximum oscillatory peak amplitude (**Supplemental Fig. S5B**, p=0.1). Interestingly, the oscillatory response duration of Oxo-M induced calcium responses increased following STIM2 KD (p<0.01) and 2-APB treatment (p<0.0001) compared to control cells (**Supplemental Fig. S5C**). In summary, these results suggest that STIM1 is required for Gα_q_-induced SOCE while STIM2 specifically modulates the frequency of Ca^2+^ oscillations following ligand-mediated SOCE.

### STIM2 is required for human oligodendrocyte progenitor differentiation

To determine whether STIM-dependent calcium oscillations may directly contribute to muscarinic M_1/3_R-mediated inhibition of hOPC differentiation, we assessed the independent effects of STIM1 and STIM2 siRNA treatment and their interactions with Oxo-M treatment on hOPC differentiation *in vitro* by three-way ANOVA (**Fig. 3A**). As shown previously^15^, Oxo-M treatment significantly blocked differentiation into O4^+^ immature OLs (F(1,3)=14.31, p=0.032, n=4 human samples). Both siRNA control and STIM1 siRNA transfected hOPCs showed reduced differentiation in response to Oxo-M (**Fig. 3B;** Holm-Sidak’s post-hoc test, control: p = 0.006; STIM1: p=0.015). Interestingly, STIM2 siRNA transfection alone or in combination with STIM1 abolished the anti-differentiative effects of Oxo-M (p > 0.2). This is consistent with a role of STIM2 downstream of Oxo-M mediated signaling.

**Figure 3.**
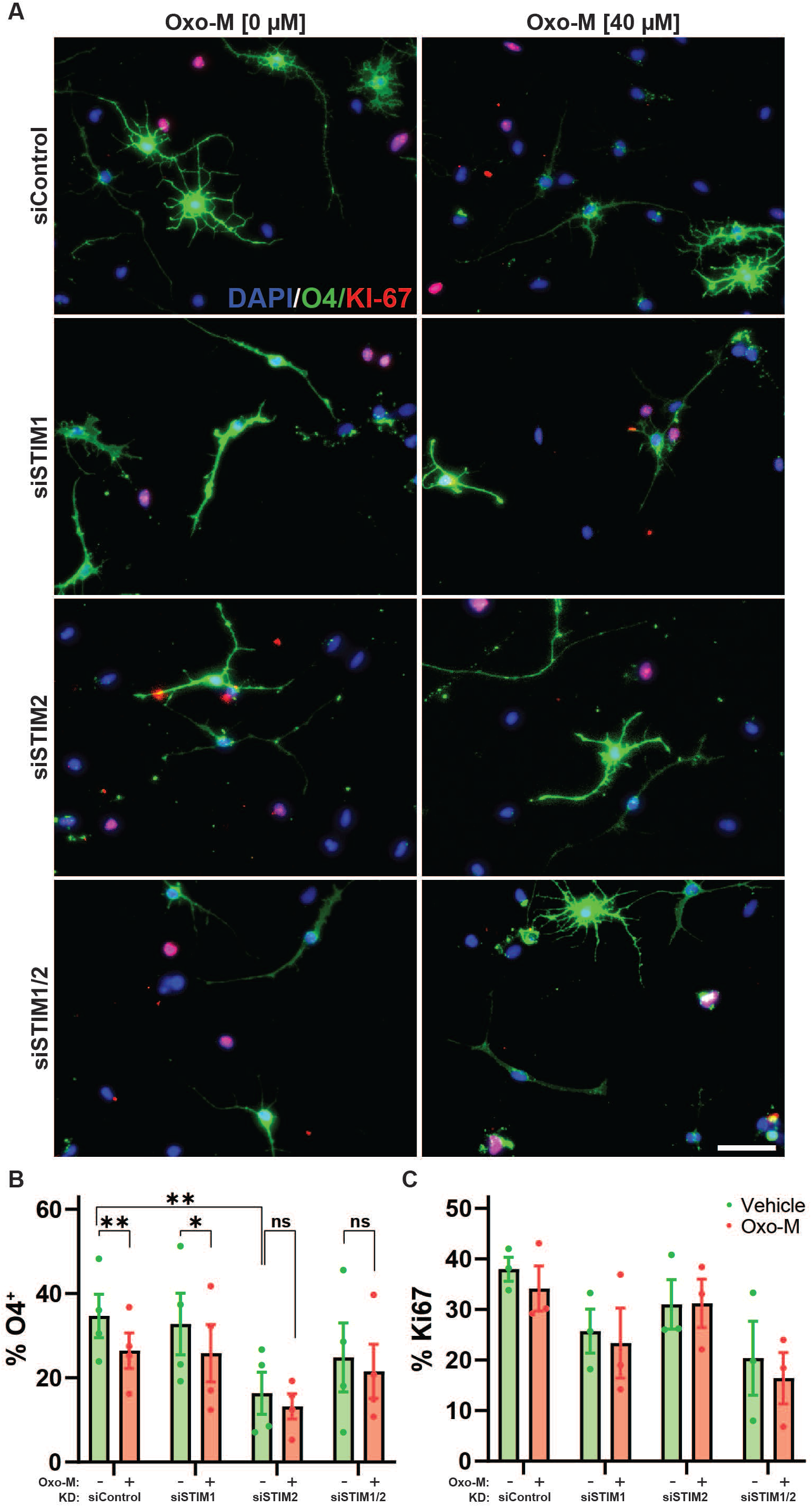
Calcium sensor STIM2 regulates human OPC differentiation and is required for the anti-differentiative effect of muscarinic agonists. hOPCs were transfected with STIM1 and/or STIM2 siRNA, or control non-targeting siRNA, and 24 h later PDGF-AA and NT-3 were removed to induce oligodendrocyte differentiation. **A**, representative images of immature O4^+^ oligodendrocytes (green) in the presence or absence of muscarinic agonist, Oxo-M [40µM]. **B**, quantification of the percentage of O4^+^ oligodendrocyte differentiation among DAPI^+^ cells (blue). Mean ± SEM shown (n=4 human fetal sample culture). Differences in O4% were analyzed by RM three-way ANOVA using OxoM, STIM1, and STIM2 as factors and repeated measures across individual patient samples. Both Oxo-M and STIM2 siRNA treatment resulted in a significant effect on O4^+^ oligodendrocyte differentiation (OxoM: F (1, 3) = 14.31, p = 0.032 and STIM2: F (1, 3) = 28.64, p = 0.0128). * p<0.05 and ** p<0.01 indicate pairwise Holm-Sidak post-hoc tests. **C**, the percentage of dividing OPCs defined as KI-67^+^/DAPI^+^ cells following siRNA transfection was determined in each group (n = 3 human samples). Neither STIM1/2 siRNA nor Oxo-M treatment significantly influenced OPC proliferation (RM three-way ANOVA, main effects p > 0.05). Scale: 50 µm.

In addition to Oxo-M dependent effects, STIM2 siRNA alone caused a significant reduction in O4^+^ OL differentiation (RM three-way ANOVA, STIM2 main effect F(1,3) = 28.64, p = 0.013, n=4 human samples) while STIM1 siRNA had no effect (p = 0.42). In the absence of Oxo-M, we observed a significant decrease in O4^+^ OL differentiation cells following STIM2 siRNA transfection relative to siControl (p=0.002). Interestingly, this effect was not observed when STIM1/2 siRNA were combined (p>0.05). These results suggested that STIM2 signaling in the absence of Gα_q_ agonist regulates hOPC differentiation and is required for the anti-differentiative effects of muscarinic agonism. We further examined the effect of STIM1/2 on morphological maturation by assessing the proportion of O4^+^ cells with simple or complex process elaboration as described ^47^ (**Supplemental Fig. S6A**). STIM1/2 KD did not influence the percentage of simple O4^+^ oligodendrocytes (3-way ANOVA, p > 0.3). Oxo-M reduced the proportion of complex O4^+^ cells following transfection with either control or STIM1 siRNA but did not influence complex O4^+^ cell number following STIM2 KD (Holm-Sidak’s post-hoc test, p < 0.05) (**Supplemental Fig. S6C**). Interestingly, in the absence of Oxo-M, STIM2 KD reduced the percentage of complex O4^+^ cells (Holm-Sidak’s post-hoc test, p =0.0013), while STIM1 KD had no effect (p=0.61).

As differentiation and OPC proliferation are intimately linked, we next examined whether ER calcium sensors STIM1/2 influenced hOPC proliferation. We quantified the proportion of dividing Ki67^+^ OPCs following transfection with STIM1/2 siRNA or control siRNA (**Fig. 3C**). Similar to previous observations ^15^, Oxo-M had no effect on hOPC proliferation. STIM1 or STIM2 siRNA treatment did not significantly influence proliferation (n = 3 human samples; three-way ANOVA, STIM1 F(1, 2) = 12.57, p = 0.071, STIM2 F (1, 2) = 6.27, p = 0.12). Taken together, these data suggest that STIM2 appears to be necessary for OL differentiation and morphological maturation and may be required for the effects of muscarinic agonist-mediated inhibition.

### Optogenetically-induced SOCE blocks hOPC differentiation and promotes proliferation

Our previous results suggest that STIM1/2 mediated SOCE may be required for sustained calcium release following Gα_q_ stimulation and thereby contribute to muscarinic inhibition of differentiation. We therefore sought to test whether oscillatory calcium SOCE directly regulates hOPC development. To this end, we expressed an optogenetically controllable STIM protein in hOPCs to enable blue-light control of SOCE in the absence of Gα_q_-ligand stimulation ^37^. Following infection with lentiviral OptoSTIM, hOPCs were transduced with lentiviral jRCaMP1a, a red-shifted calcium indicator, to simultaneously measure intracellular Ca^2+^ while induced SOCE ^35^. Blue light activation of OptoSTIM resulted in a consistent and robust oscillatory calcium influx which could be sustained for several hours in a frequency-specific manner (**Fig. 4A-B**). To avoid summation of OptoSTIM-induced Ca^2+^ release and ensure a distinct oscillatory wave form, based on the dissociation kinetics of OptoSTIM ^37^, we selected an upper frequency of 3.33 mHz to model that of 10mHz waves observed following muscarinic receptor activation. As a control, we selected a lower frequency of 0.55 mHz which is below the lower threshold associated with frequency-dependent decoding in other non-excitable cells ^28,48^. We matched the total blue-light exposure over time between frequencies by pulsing for 330 and 2000 ms seconds at 3.33 and 0.55 mHz respectively. Based on jRCaMP1a imaging, oscillatory SOCE could be maintained for at least three hours in hOPCs *in vitro* and was clearly observable at 18 hours with 2000 ms pulses without loss of signal (**Fig. 4B**, n = 3 human samples). Importantly, we found that the total calcium release defined by AUC over the 2.5 h period was equivalent between groups (**Fig. 4C**; RM one-way ANOVA, Holm-Sidak’s post-hoc test, p = 0.49). As such, we could assess frequency dependent SOCE effects while controlling for total calcium flux.

**Figure 4.**
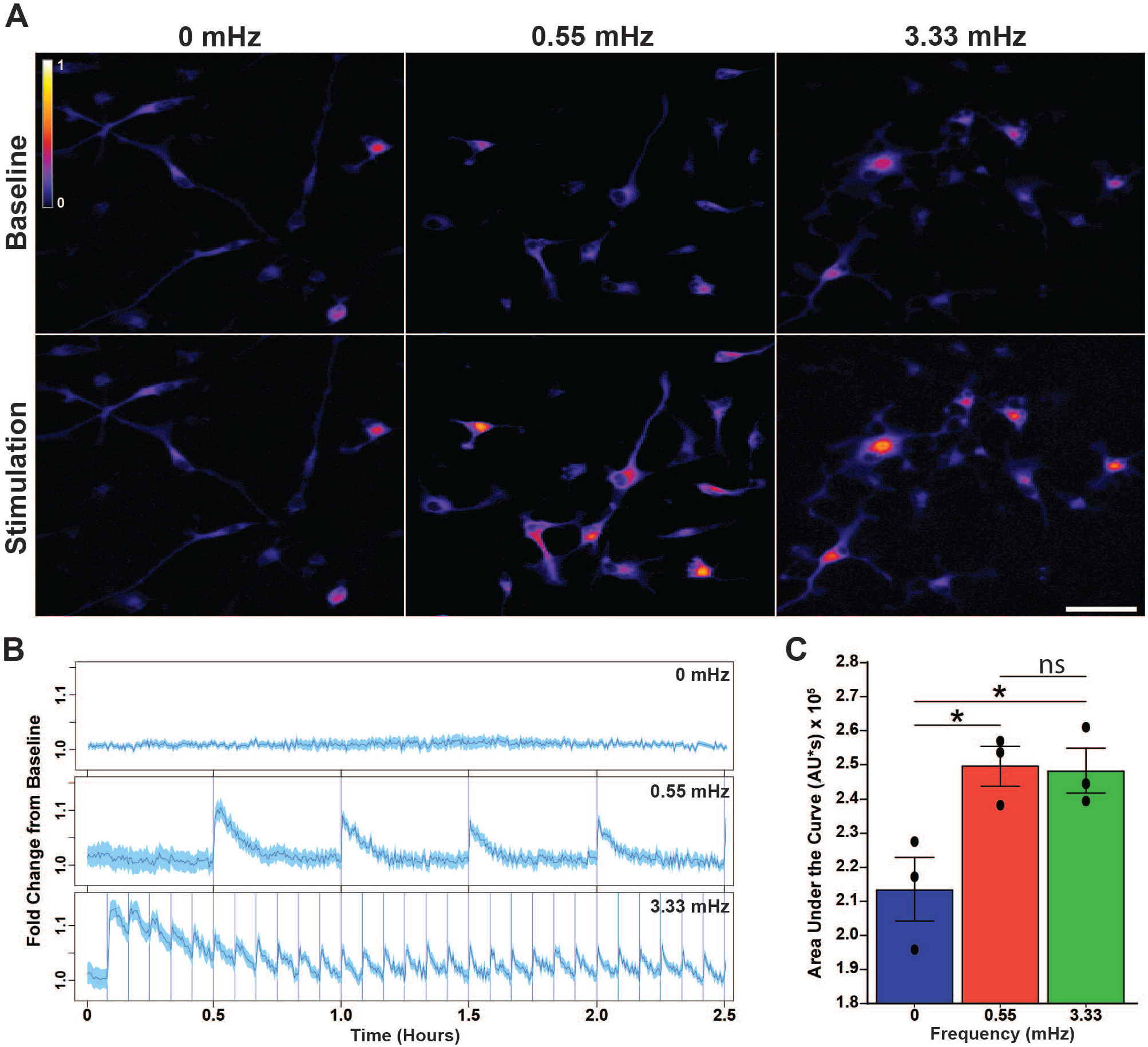
OptoSTIM drives oscillatory calcium responses in hOPCs in the absence of Gα_q_-coupled signaling. hOPCs were infected with OptoSTIM and jRCaMP1a lentiviruses to induce SOCE and observe calcium responses simultaneously. Cells were cultured in a temperature/CO_2_ controlled chamber during time-lapse fluorescence microscopy. Blue light (488nm) excitation was used to induce SOCE at distinct frequencies using 5 minute (3.33 mHz) or 30 minute (0.55 mHz) stimulation intervals respectively, Blue light exposures were matched to ensure an equivalent amount of blue light stimulation across conditions (330 and 2000 ms per stimulus, respectively). Changes in intracellular Ca^2+^ concentration were assessed by jRCaMP1a fluorescence every 20 seconds. **A**, representative integrated calcium response to OptoSTIM-induced SOCE in the absence of Gα_q_-receptor activation. Images shown represent total fluorescence across 2.5 hours of stimulation following a 30-minute baseline period. **B**, jRCaMP1a traces of mean ± SEM (blue shading) normalized calcium responses to 0.55 mHz and 3.33 mHz frequencies over the course of 2.5 hours following a 30m baseline with no stimulation (n≥50 cells from a single representative biological replicate). SOCE was robustly induced throughout the imaging period and directly corresponded with the timing of blue light stimulation (vertical blue lines in middle and lower traces). **C**, quantification of integrated calcium responses at different stimulation frequencies measured for the duration of the stimulation period. Mean ± SEM of results from three independent experiments (n=3 independent human fetal sample preparations) with >50 cells analyzed per condition/ replicate. * p<0.05, RM one-way ANOVA with Holm-Sidak’s post-hoc test. Scale: 50 µm.

We examined the effect of optogenetically-induced oscillatory SOCE on hOPC proliferation and differentiation (**Fig. 5**). Using time lapse microscopy and concurrent intermittent blue-light stimulation to induce SOCE, we assessed the number of mitotic events (**Fig. 5A**) that occurred in the presence of both high and low frequency OptoSTIM1-induced SOCE. Induction of high frequency (3.33 mHz) oscillatory calcium entry produced a clear increase in the number of cumulative mitotic events at 24h and 36h post stimulation respectively (**Fig. 5B**, quantified in **C**; RM one-way ANOVA with Holm-Sidak’s post-hoc test, p = 0.006, n = 4 human samples). After 36 hours of stimulation, high frequency SOCE resulted in a 40% increase rate of cell division relative to unstimulated control hOPCs (3.33 mHz: 25.6 ± 4.9% vs. control: 19.1 ± 4.7%, p = 0.014) (**Fig. 5C**). In contrast, low frequency SOCE had no effect on proliferation (0.55 mHz; 17.6 ± 5.9%, p = 0.38). Next, we investigated the effects of long term OptoSTIM induction on hOPC differentiation (**Fig. 5D**). We observed no gross alterations in transduced cell number following blue-light stimulation (**Fig. 5E**; RM one-way ANOVA with Holm-Sidak’s post-hoc test, p = 0.90, n = 4 human samples). Importantly, induction of high frequency SOCE substantially impaired hOPC differentiation by 50% relative to unstimulated control hOPCs (**Fig. 5F**; 3.33 mHz: 7.5 ± 1.7% O4^+^/GFP^+^ cells vs. control: 16.6 ± 3.4, p = 0.02). Differentiation by hOPCs which received induction of low frequency SOCE was not significantly different from unstimulated control cells (0.55 mHz: 12.83 ± 3.3%). Moreover, there was a significant reduction in the number of O4^+^ cells with complex branch morphology (>3 processes) following 3.33 mHz stimulation compared to control cells (**Fig. 5G;** RM one-way ANOVA with Holm-Sidak’s post-hoc test, p = 0.04). This suggests that induction of oscillatory SOCE was sufficient to inhibit hOPC morphological maturatio. Collectively these results indicate that sustained high frequency oscillatory SOCE was sufficient to both block hOPC differentiation and augment hOPC proliferation.

**Figure 5.**
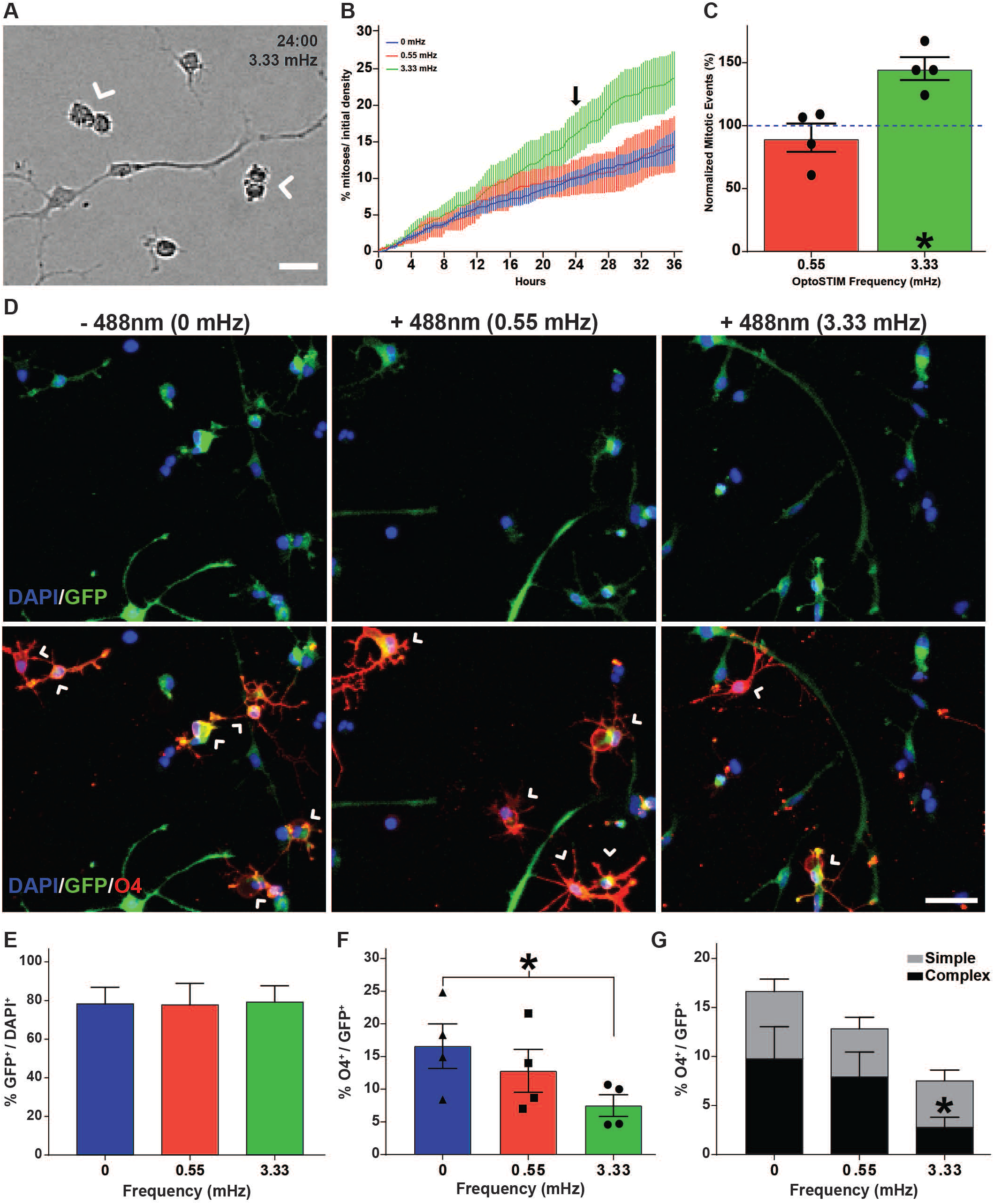
SOCE is sufficient to block hOPC differentiation and promote hOPC mitosis. Matched cultures of primary hOPCs were infected with OptoSTIM lentivirus and imaged for 2-3 days in an on-stage temperature/CO_2_ controlled chamber. Bright field images were acquired every 5 minutes. OptoSTIM-mediated SOCE was induced with intermittent 488nm excitation (5m or 30m interval for 330ms and 2000ms duration, respectively). **A**, hOPC proliferation was quantified by counting mitotic events (white arrowheads). **B**, the cumulative proportion of cells undergoing mitosis was quantified over 36 h following the onset of blue light stimulation. **C**, proliferation rates were normalized to the matched unstimulated condition. High frequency (3.33 mHz) OptoSTIM induction of SOCE resulted in a significant increase in hOPC mitotic events compared to control unstimulated and low frequency (0.55 mHz) stimulated hOPCs (n=4 independent human fetal sample culture preparations). **D**, the effect of OptoSTIM-induced SOCE on hOPC differentiation was determined by O4 immunocytochemistry (red) at 48 hrs post-stimulation. OptoSTIM transduced cells were identified by GFP expression which was uniform across experimental conditions (**E**). **F**, quantification of O4^+^ oligodendrocyte differentiation following OptoSTIM-induced SOCE (% O4^+^/GFP^+^). **G**, analysis of O4^+^ morphological maturation. High frequency (3.33 mHz) SOCE induced a significant reduction in O4^+^ cells as well as a significant reduction in complex branching morphology relative to control hOPCs. * p<0.05, Holm-Sidak’s post-hoc test. Mean ± SEM shown for each panel (n=4 independent human fetal samples). Scale: 25 µm.

## DISCUSSION

In the present study, we demonstrate that a subset of Gα_q_-receptors expressed by human primary OPCs, including muscarinic M_1_ and M_3_ receptors (M_1/3_R) and metabotropic-glutamate-receptor-5 (mGluR5), induced prolonged oscillatory calcium signaling and that the ability to induce oscillatory calcium signaling correlated with their capacity to block differentiation. We found that a shared mechanism downstream muscarinic Gα_q_-receptor induced oscillatory signaling was the activation of store operated calcium entry (SOCE). Inhibition with SOCE antagonists or knockdown of ER-localized SOCE calcium sensors STIM1/2 reduced the magnitude of calcium signaling induced by muscarinic agonist stimulation. STIM2 ablation selectively reduced calcium oscillation frequency and spontaneous OL differentiation *in vitro* and, unlike control OPCs, treatment with muscarinic ligand did not further reduce differentiation following STIM2 knockdown. While SOCE exerted a major influence on calcium oscillations, as these experiments were performed using constant Gα_q_-receptor agonist exposure, we cannot exclude the requirement of IP_3_-mediated release for the maintenance of prolonged calcium oscillations. Together, these data suggested a functional linkage between muscarinic receptor signaling and SOCE. Using an optogenetic approach to selectively activate SOCE in OPCs, we found that SOCE was sufficient to modulate OL differentiation in the absence of Gα_q_ dependent signaling suggesting that SOCE is a key facet that impairs OL differentiation and remyelination. Furthermore, the frequency of SOCE rather than the overall amount of calcium influx due to SOCE was found to be vital in mediating the effect of calcium entry on differentiation and proliferation. This suggests that the calcium pathway includes frequency-dependent ‘decoders’ which regulate OL differentiation and that modulation of calcium frequency might represent an important mechanism by which diverse signals are integrated by parenchymal OPCs.

In drug development, G-protein coupled receptors (GPCRs) are attractive candidates for therapeutic intervention and recently many small molecules that target GPCRs have been identified that modulate OPC function and OL differentiation to promote myelin repair ^49^. For example, GPR17 which largely couples via Gα_i/o_ was found to negatively regulate OPC differentiation while knockout was observed to promote remyelination ^50^. Muscarinic receptor antagonists have displayed a remarkable capacity to improve OL differentiation and remyelination in various animal models. Using genetic approaches, the effects of these drugs have been localized to M_1_R and M_3_R. Both of these receptors represent canonical Gα_q_-coupled receptors. Indeed, when Gα coupling was systematically analyzed across different subunits, M_3_R displayed exquisite selectivity for rapid activation for Gα_q_/Gα_11_ coupling ^51^. In this study, we focused on Gα_q_-coupled receptors to determine if a shared characteristic of Gα_q_ receptors could be used to predict their ability to prevent efficient OL differentiation and thereby impair remyelination. We analyzed RNA-seq analysis of primary hOPC to identity those Gα_q_-coupled receptors expressed at the mRNA level and hypothesized that receptor-specific ligand activation would produce canonical oscillatory calcium signaling and block hOPC differentiation. Surprisingly, only a small subset of pharmacological Gα_q_ ligands were capable of inducing prolonged calcium oscillation in hOPCs. Intriguingly, this correlated with the ability of ligands to effectively block OL differentiation. CHPG, which activates mGluR_5_ in a highly selective manner, was the only other ligand capable of inducing calcium oscillations in a large fraction of hOPCs. The specificity of Gα_q_-receptor mediated signaling is consistent with previous results which have shown that while endothelin (ET) agonist treatment induces a monotonic calcium response ^24^ while metabotropic glutamate receptor agonists induce calcium oscillations ^19^. mGluR_5_ couples primarily via Gα_q_ and does not have appreciable activity at Gα_i_. In a similar manner to M_3_R, mGluR_5_ mRNA was relatively highly expressed in rodent OPCs and down-regulated following differentiation ^19^. Herein, we found that treatment with CHPG resulted in a dose-dependent blockade of hOPC differentiation.

In contrast, the other Gα_q_ couple receptors that elicited calcium responses in OPCs induced calcium release that was either considerably shorter in duration and/or were largely monotonic in nature. These differences could be due to differences in overall level of expression and/or differences in receptor coupling. From a large-scale screen of G-protein coupling ^52^, we observed that many of these receptors were promiscuously coupled with other Gα subunits. For example, sphingosine-1-phosphate-receptor-1 (S1PR1) was observed as primarily coupled via Gα_i_ and Gα_o_ with little evidence of Gα_q_ coupling. This difference corresponded with the observed function difference following activation of S1PR1 which results in OL differentiation from human OPCs ^53^. Furthermore, endothelin-B receptor (ENDRB), thrombin receptor II (F2R) and purinergic receptor Y1 (P2RY1) also show promiscuous coupling with multiple Gα proteins upon activation. Together this suggests that selectively in coupling may influence the net influence on downstream signaling and the nature of the resultant calcium response. As the coupling of GPCRs to specific Gα proteins are not absolute and in many cases is promiscuous, we anticipate that biased ligands or antagonists capable of promoting or blocking coupling along specific pathways may represent optimal approaches to promote OPC differentiation.

The mechanisms which shape the calcium-response following ligand application are complex and likely dependent on multiple interacting proteins. Among these, Gα subunits form specific complexes with their cognate Gβγ proteins which possess their own signaling capability and modulate the signaling of Gα subunits they interact with ^54^. Among these, specific isoforms of phosphoinositide 3-kinase (PI3K) and adenylate cyclase, voltage-operated calcium channels (VOCC) and inwardly rectifying K^+^ channel can be regulated by Gβγ. Activation of PI3K/Akt ^55^, adenylate cyclase ^56^, and VOCC ^57^ are associated with promotion of OPC differentiation suggesting that Gβγ activation may counteract oscillatory calcium signaling. For muscarinic receptor signaling, in rat pituitary cells G_β3_ is necessary for the carbachol-mediated calcium current ^58^, while in a rat leukemia cell line G_β1_ and G_β4_ were more important in the regulation of this current ^59^. In primary hOPCs, G_β4_ ablation did not influence calcium signaling following Oxo-M treatment ^60^, though redundancy with G_β1_ and G_β2_ is possible as these subunits are expressed at substantially higher levels in human, rat, and mouse OPCs.

In addition to the G_βγ_ subunit specific regulation, receptor-specific activation of regulators of G-protein signaling (RGS) proteins provide specificity for G-protein coupling and modulate the kinetics of second messenger activation ^61^. RGS proteins promote termination of GPCR signaling via GTPase activation and prevent GPCR signaling in the absence of ligand. Following ligand activation and elevated Ca^2+^, RGS proteins exhibit a cyclical pattern of association with phosphatidylinositol-3,4,5-trisphosphate (PIP_3_) and activated calcium/calmodulin (Ca^2+^/CaM) followed by re-association with the GPCR complex after Ca^2+^ levels return to baseline. This allows a second cycle to start and initiate another calcium spike. The kinetics of these steps, especially the slow dissociation of RGS protein from Ca^2+^/CaM, contributes to the resultant frequency of calcium oscillations ^62^. While the role of specific RGS proteins in the OPC and OL lineage is poorly understood, several RGS proteins are expressed in OPCs ^46^. Among these, RGS4 has been shown to be a potent negative regulator of muscarinic receptor signaling in insulinoma cells ^63^. Also noteworthy, the R7 family of RGS proteins (RGS6/7/9/11) form heterodimers with G-protein β subunit Gβ5 (GNB5) which similarly attenuates M_3_R-dependent Ca^2+^ signaling ^64^.

Once activated, SOCE is associated with many cellular functions in neural progenitors such as proliferation ^65^ and differentiation ^66,67^. In OPCs, SOCE can modulate PDGF-AA induced OPC proliferation ^30^. However, given the known role of PI3K/Akt signaling downstream of PDGF receptor, SOCE is unlikely to be a critical component of PDGF signaling. Likewise, the cellular role of calcium oscillations in OPC soma is not yet known. Our data using optogenetic induction of SOCE strongly suggest that oscillatory SOCE alone is sufficient to modulate both OPC proliferation and differentiation. While we were limited by the dissociation kinetics of OptoSTIM to an upper limit of 3.33 mHz stimulation, this frequency fits within the known frequencies of calcium oscillations in non-excitable cells and may recruit NFAT, NFκB and other downstream pathways ^48^. By matching total calcium influx during the period of exposure, we were able to demonstrate frequency dependence as a substantially lower frequency (0.55 mHz) was unable to modulate proliferation or differentiation. New approaches will be required to determine whether higher frequency SOCE can further alter these critical cellular processes.

SOCE itself is dependent on the association of CRAC subunit comprising STIM ER-calcium sensors and ORAI channel subunits. While STIM1 and STIM2 contributed to different aspects of muscarinic-induced SOCE and sustained oscillatory calcium signaling in hOPCs, we found that only STIM2 knockdown reduced spontaneous OL differentiation. Previous reports have found that apart from its role in agonist mediated SOCE, STIM2 signaling often persists to a lesser degree in the absence of an ER-release stimulus. STIM2 responds to small fluctuations in ER calcium levels, and drives small local SOCE calcium entry when local calcium levels fall below baseline (reviewed in ^68^). It is also known that STIM2 can regulate endogenous differentiation of other cell types through modulation of cytoplasmic calcium levels following STIM1-induced oscillatory SOCE ^69^. Furthermore, small spontaneous SOCE calcium transients in the absence of agonist have been observed in mouse OPCs, and have been implicated in the regulation of differentiation ^70^. It is possible, therefore, that STIM2 is required for these spontaneous transients and thus STIM2 KD may inhibit differentiation. In contrast, our data is consistent with prolonged oscillatory SOCE, either produced optogenetically or in response to M_3_R and mGluR_5_ agonists, causing a blockade of hOPC differentiation via global calcium-mediated changes. Additional studies will be necessary to tease apart the specific mechanisms by which STIM2 directly influences oligodendrocyte differentiation.

The ER-calcium sensors STIM1/2 physically and functionally interact with one another to regulate SOCE ^68^. The relative abundance of STIM1 and STIM2 vary greatly between tissues and individual cell types. In human OPCs, STIM2 is more than 3-fold more abundant than STIM1 mRNA but both genes are relatively highly expressed (>10 FPKM). In contrast, both STIM1 and STIM2 are expressed at equivalent levels in mouse OPCs ^46^. As the differences in the relative expression of STIM1/2 influences the resultant properties of SOCE in response to store depletion (reviewed in ^71^), the differences in STIM1:STIM2 ratio in mouse and human may therefore underlie some of the species-specific functional differences in OPC behavior ^72^. Both STIM1 and STIM2 have been shown to contribute to oscillatory SOCE in a context dependent manner ^73,74^. Our data support a specific role of STIM2 in the regulation of OPC calcium oscillations and suggest that expression of STIM2 is required for muscarinic receptor induced differentiation blockade in hOPCs. Together with the direct regulation of OPC differentiation via optogenetically-gated SOCE, these data suggest that SOCE mediated via STIM activation will likely play an important role in mediating the negative effects of Gα_q_-coupled receptors capable of blocking oligodendrocyte differentiation in health and disease.

## Acknowledgements

This work was supported by grants from the NINDS (R01NS104021), the National Multiple Sclerosis Society (RG 5505-A-2, RG 5110A1/1), the Kalec Multiple Sclerosis Foundation, the Change MS Foundation, and the Skarlow Memorial Trust. JJP received support from NIGMS (R25GM09545902), NCATS (UL1TR001412-S1). JJP and RAS received support from NYSTEM Stem Cells in Regenerative Medicine Fellowship (#C302090G). In addition, we would like to acknowledge the assistance of the Confocal Microscope and Flow Cytometry Facility in the School of Medicine and Biomedical Sciences, University at Buffalo and thank Dr. Pablo Paez for advice with calcium imaging of OPCs.

## Notes

### Competing Interest Statement

The authors have declared no competing interest.

## REFERENCES

1 Franklin, R. J. M. & Ffrench-Constant, C. Regenerating CNS myelin - from mechanisms to experimental medicines. Nature reviews. Neuroscience 18, 753–769, doi:10.1038/nrn.2017.136 (2017).

2 Faissner, S., Plemel, J. R., Gold, R. & Yong, V. W. Progressive multiple sclerosis: from pathophysiology to therapeutic strategies. Nat Rev Drug Discov 18, 905–922, doi:10.1038/s41573-019-0035-2 (2019).

3 Kuhlmann, T. et al. Differentiation block of oligodendroglial progenitor cells as a cause for remyelination failure in chronic multiple sclerosis. Brain : a journal of neurology 131, 1749–1758, doi:10.1093/brain/awn096 (2008).

4 Wolswijk, G. Chronic stage multiple sclerosis lesions contain a relatively quiescent population of oligodendrocyte precursor cells. J Neurosci 18, 601–609 (1998).

5 Thornton, M. A. & Hughes, E. G. Neuron-oligodendroglia interactions: Activity-dependent regulation of cellular signaling. Neuroscience letters 727, 134916, doi:10.1016/j.neulet.2020.134916 (2020).

6 Ortiz, F. C. et al. Neuronal activity in vivo enhances functional myelin repair. JCI Insight 5, doi:10.1172/jci.insight.123434 (2019).

7 Gautier, H. O. et al. Neuronal activity regulates remyelination via glutamate signalling to oligodendrocyte progenitors. Nat Commun 6, 8518, doi:10.1038/ncomms9518 (2015).

8 Bernstein, M., Lyons, S. A., Moller, T. & Kettenmann, H. Receptor-mediated calcium signalling in glial cells from mouse corpus callosum slices. Journal of neuroscience research 46, 152–163, doi:10.1002/(SICI)1097-4547(19961015)46:2<152::AID-JNR3>3.0.CO;2-G (1996).

9 Paez, P. M. & Lyons, D. A. Calcium Signaling in the Oligodendrocyte Lineage: Regulators and Consequences. Annual review of neuroscience 43, 163–186, doi:10.1146/annurev-neuro-100719-093305 (2020).

10 Berridge, M. J. & Irvine, R. F. Inositol phosphates and cell signalling. Nature 341, 197–205, doi:10.1038/341197a0 (1989).

11 Mei, F. et al. Accelerated remyelination during inflammatory demyelination prevents axonal loss and improves functional recovery. Elife 5, e18246, doi:10.7554/eLife.18246 (2016).

12 Welliver, R. R. et al. Muscarinic Receptor M3R Signaling Prevents Efficient Remyelination by Human and Mouse Oligodendrocyte Progenitor Cells. J Neurosci 38, 6921–6932, doi:10.1523/JNEUROSCI.1862-17.2018 (2018).

13 Mei, F. et al. Micropillar arrays as a high-throughput screening platform for therapeutics in multiple sclerosis. Nature medicine 20, 954–960, doi:10.1038/nm.3618 (2014).

14 Deshmukh, V. A. et al. A regenerative approach to the treatment of multiple sclerosis. Nature 502, 327–332, doi:10.1038/nature12647 (2013).

15 Abiraman, K. et al. Anti-muscarinic adjunct therapy accelerates functional human oligodendrocyte repair. J Neurosci 35, 3676–3688, doi:10.1523/JNEUROSCI.3510-14.2015 (2015).

16 Gadea, A., Aguirre, A., Haydar, T. F. & Gallo, V. Endothelin-1 regulates oligodendrocyte development. J Neurosci 29, 10047–10062, doi:10.1523/JNEUROSCI.0822-09.2009 (2009).

17 Agresti, C. et al. ATP regulates oligodendrocyte progenitor migration, proliferation, and differentiation: involvement of metabotropic P2 receptors. Brain Res Brain Res Rev 48, 157–165, doi:10.1016/j.brainresrev.2004.12.005 (2005).

18 Takeda, M., Nelson, D. J. & Soliven, B. Calcium signaling in cultured rat oligodendrocytes. Glia 14, 225–236, doi:10.1002/glia.440140308 (1995).

19 Luyt, K., Varadi, A., Durant, C. F. & Molnar, E. Oligodendroglial metabotropic glutamate receptors are developmentally regulated and involved in the prevention of apoptosis. J Neurochem 99, 641–656, doi:10.1111/j.1471-4159.2006.04103.x (2006).

20 Fan, L. W. et al. Exposure to serotonin adversely affects oligodendrocyte development and myelination in vitro. J Neurochem 133, 532–543, doi:10.1111/jnc.12988 (2015).

21 Jung, C. G. et al. Functional consequences of S1P receptor modulation in rat oligodendroglial lineage cells. Glia 55, 1656–1667, doi:10.1002/glia.20576 (2007).

22 Chattopadhyay, N. et al. Calcium receptor expression and function in oligodendrocyte commitment and lineage progression: potential impact on reduced myelin basic protein in CaR-null mice. Journal of neuroscience research 86, 2159–2167, doi:10.1002/jnr.21662 (2008).

23 Coppi, E. et al. Adenosine A2B receptors inhibit K(+) currents and cell differentiation in cultured oligodendrocyte precursor cells and modulate sphingosine-1-phosphate signaling pathway. Biochem Pharmacol 177, 113956, doi:10.1016/j.bcp.2020.113956 (2020).

24 Jung, K. J. et al. The role of endothelin receptor A during myelination of developing oligodendrocytes. J Korean Med Sci 26, 92–99, doi:10.3346/jkms.2011.26.1.92 (2011).

25 Yuen, T. J. et al. Identification of endothelin 2 as an inflammatory factor that promotes central nervous system remyelination. Brain : a journal of neurology 136, 1035–1047, doi:10.1093/brain/awt024 (2013).

26 Dukala, D. E. & Soliven, B. S1P1 deletion in oligodendroglial lineage cells: Effect on differentiation and myelination. Glia 64, 570–582, doi:10.1002/glia.22949 (2016).

27 Prakriya, M. & Lewis, R. S. Store-Operated Calcium Channels. Physiol Rev 95, 1383–1436, doi:10.1152/physrev.00020.2014 (2015).

28 Smedler, E. & Uhlen, P. Frequency decoding of calcium oscillations. Biochim Biophys Acta 1840, 964–969, doi:10.1016/j.bbagen.2013.11.015 (2014).

29 Samanta, K. & Parekh, A. B. Spatial Ca(2+) profiling: decrypting the universal cytosolic Ca(2+) oscillation. J Physiol 595, 3053–3062, doi:10.1113/JP272860 (2017).

30 Paez, P. M. et al. Regulation of store-operated and voltage-operated Ca2+ channels in the proliferation and death of oligodendrocyte precursor cells by golli proteins. ASN Neuro 1, doi:10.1042/AN20090003 (2009).

31 Schmidt, C., Ohlemeyer, C., Kettenmann, H., Reutter, W. & Horstkorte, R. Incorporation of N-propanoylneuraminic acid leads to calcium oscillations in oligodendrocytes upon the application of GABA. Febs Letters 478, 276–280, doi:Doi 10.1016/S0014-5793(00)01868-8 (2000).

32 Paez, P. M., Fulton, D., Spreuer, V., Handley, V. & Campagnoni, A. T. Modulation of canonical transient receptor potential channel 1 in the proliferation of oligodendrocyte precursor cells by the golli products of the myelin basic protein gene. J Neurosci 31, 3625–3637, doi:10.1523/JNEUROSCI.4424-10.2011 (2011).

33 Conway, G. D., O’Bara, M. A., Vedia, B. H., Pol, S. U. & Sim, F. J. Histone deacetylase activity is required for human oligodendrocyte progenitor differentiation. Glia 60, 1944–1953, doi:10.1002/glia.22410 (2012).

34 Pol, S. U. et al. Sox10-MCS5 enhancer dynamically tracks human oligodendrocyte progenitor fate. Experimental neurology 247, 694–702, doi:10.1016/j.expneurol.2013.03.010 (2013).

35 Dana, H. et al. Sensitive red protein calcium indicators for imaging neural activity. Elife 5, doi:10.7554/eLife.12727 (2016).

36 Sevin, C. et al. Intracerebral adeno-associated virus-mediated gene transfer in rapidly progressive forms of metachromatic leukodystrophy. Hum Mol Genet 15, 53–64, doi:10.1093/hmg/ddi425 (2006).

37 Kyung, T. et al. Optogenetic control of endogenous Ca(2+) channels in vivo. Nature biotechnology 33, 1092–1096, doi:10.1038/nbt.3350 (2015).

38 Wang, J. et al. Paired Related Homeobox Protein 1 Regulates Quiescence in Human Oligodendrocyte Progenitors. Cell reports 25, 3435–3450 e3436, doi:10.1016/j.celrep.2018.11.068 (2018).

39 Geraerts, M., Willems, S., Baekelandt, V., Debyser, Z. & Gijsbers, R. Comparison of lentiviral vector titration methods. BMC biotechnology 6, 34, doi:10.1186/1472-6750-6-34 (2006).

40 Edelstein, A., Amodaj, N., Hoover, K., Vale, R. & Stuurman, N. Computer control of microscopes using microManager. Current protocols in molecular biology Chapter 14, Unit14.20, doi:10.1002/0471142727.mb1420s92 (2010).

41 Stockley, J. H. et al. Surpassing light-induced cell damage in vitro with novel cell culture media. Sci Rep 7, 849, doi:10.1038/s41598-017-00829-x (2017).

42 Tinevez, J. Y. et al. TrackMate: An open and extensible platform for single-particle tracking. Methods 115, 80–90, doi:10.1016/j.ymeth.2016.09.016 (2017).

43 Treiman, M., Caspersen, C. & Christensen, S. B. A tool coming of age: thapsigargin as an inhibitor of sarco-endoplasmic reticulum Ca(2+)-ATPases. Trends in pharmacological sciences 19, 131–135, doi:10.1016/s0165-6147(98)01184-5 (1998).

44 DeHaven, W. I., Smyth, J. T., Boyles, R. R., Bird, G. S. & Putney, J. W., Jr. Complex actions of 2-aminoethyldiphenyl borate on store-operated calcium entry. The Journal of biological chemistry 283, 19265–19273, doi:10.1074/jbc.M801535200 (2008).

45 Harper, J. L. et al. Dihydropyridines as inhibitors of capacitative calcium entry in leukemic HL-60 cells. Biochem Pharmacol 65, 329–338, doi:10.1016/s0006-2952(02)01488-0 (2003).

46 Zhang, Y. et al. An RNA-sequencing transcriptome and splicing database of glia, neurons, and vascular cells of the cerebral cortex. J Neurosci 34, 11929–11947, doi:10.1523/JNEUROSCI.1860-14.2014 (2014).

47 Huang, J. K. et al. Retinoid × receptor gamma signaling accelerates CNS remyelination. Nat Neurosci 14, 45–53, doi:10.1038/nn.2702 (2011).

48 Boulware, M. J. & Marchant, J. S. Timing in cellular Ca2+ signaling. Curr Biol 18, R769–R776, doi:10.1016/j.cub.2008.07.018 (2008).

49 Mogha, A., D’Rozario, M. & Monk, K. R. G Protein-Coupled Receptors in Myelinating Glia. Trends in pharmacological sciences 37, 977–987, doi:10.1016/j.tips.2016.09.002 (2016).

50 Ou, Z. et al. Olig2-Targeted G-Protein-Coupled Receptor Gpr17 Regulates Oligodendrocyte Survival in Response to Lysolecithin-Induced Demyelination. J Neurosci 36, 10560–10573, doi:10.1523/JNEUROSCI.0898-16.2016 (2016).

51 Masuho, I. et al. Distinct profiles of functional discrimination among G proteins determine the actions of G protein-coupled receptors. Sci Signal 8, ra123, doi:10.1126/scisignal.aab4068 (2015).

52 Inoue, A. et al. Illuminating G-Protein-Coupling Selectivity of GPCRs. Cell 177, 1933–1947 e1925, doi:10.1016/j.cell.2019.04.044 (2019).

53 Cui, Q. L., Fang, J., Kennedy, T. E., Almazan, G. & Antel, J. P. Role of p38MAPK in S1P receptor-mediated differentiation of human oligodendrocyte progenitors. Glia 62, 1361–1375, doi:https://doi.org/10.1002/glia.22688 (2014).

54 Khan, S. M. et al. The expanding roles of Gbetagamma subunits in G protein-coupled receptor signaling and drug action. Pharmacological reviews 65, 545–577, doi:10.1124/pr.111.005603 (2013).

55 Goebbels, S. et al. Elevated phosphatidylinositol 3,4,5-trisphosphate in glia triggers cell-autonomous membrane wrapping and myelination. J Neurosci 30, 8953–8964, doi:10.1523/jneurosci.0219-10.2010 (2010).

56 Raible, D. W. & McMorris, F. A. Induction of oligodendrocyte differentiation by activators of adenylate cyclase. Journal of neuroscience research 27, 43–46, doi:https://doi.org/10.1002/jnr.490270107 (1990).

57 Cheli, V. T. et al. Conditional Deletion of the L-Type Calcium Channel Cav1.2 in Oligodendrocyte Progenitor Cells Affects Postnatal Myelination in Mice. J Neurosci 36, 10853–10869, doi:10.1523/JNEUROSCI.1770-16.2016 (2016).

58 Kleuss, C., Scherübl, H., Hescheler, J., Schultz, G. & Wittig, B. Different β-subunits determine G-protein interaction with transmembrane receptors. Nature 358, 424–426, doi:10.1038/358424a0 (1992).

59 Dippel, E., Kalkbrenner, F., Wittig, B. & Schultz, G. A heterotrimeric G protein complex couples the muscarinic m1 receptor to phospholipase C-beta. Proceedings of the National Academy of Sciences 93, 1391–1396, doi:10.1073/pnas.93.4.1391 (1996).

60 Pol, S. U. et al. Network-Based Genomic Analysis of Human Oligodendrocyte Progenitor Differentiation. Stem cell reports 9, 710–723, doi:10.1016/j.stemcr.2017.07.007 (2017).

61 Kach, J., Sethakorn, N. & Dulin, N. O. A finer tuning of G-protein signaling through regulated control of RGS proteins. Am J Physiol Heart Circ Physiol 303, H19–35, doi:10.1152/ajpheart.00764.2011 (2012).

62 Luo, X., Popov, S., Bera, A. K., Wilkie, T. M. & Muallem, S. RGS Proteins Provide Biochemical Control of Agonist-Evoked [Ca2+]i Oscillations. Molecular cell 7, 651–660, doi:https://doi.org/10.1016/S1097-2765(01)00211-8 (2001).

63 Ruiz de Azua, I. et al. RGS4 is a negative regulator of insulin release from pancreatic β-cells in vitro and in vivo. Proceedings of the National Academy of Sciences 107, 7999–8004, doi:10.1073/pnas.1003655107 (2010).

64 Karpinsky-Semper, D., Volmar, C. H., Brothers, S. P. & Slepak, V. Z. Differential effects of the Gβ5-RGS7 complex on muscarinic M3 receptor-induced Ca2+ influx and release. Mol Pharmacol 85, 758–768, doi:10.1124/mol.114.091843 (2014).

65 Domenichini, F. et al. Store-Operated Calcium Entries Control Neural Stem Cell Self-Renewal in the Adult Brain Subventricular Zone. Stem Cells 36, 761–774, doi:10.1002/stem.2786 (2018).

66 Forostyak, O. et al. Physiology of Ca(2+) signalling in stem cells of different origins and differentiation stages. Cell Calcium 59, 57–66, doi:10.1016/j.ceca.2016.02.001 (2016).

67 Gopurappilly, R., Deb, B. K., Chakraborty, P. & Hasan, G. Stable STIM1 Knockdown in Self-Renewing Human Neural Precursors Promotes Premature Neural Differentiation. Front Mol Neurosci 11, 178, doi:10.3389/fnmol.2018.00178 (2018).

68 López, E., Salido, G. M., Rosado, J. A. & Berna-Erro, A. Unraveling STIM2 function. J Physiol Biochem 68, 619–633, doi:10.1007/s13105-012-0163-1 (2012).

69 Darbellay, B. et al. Human muscle economy myoblast differentiation and excitation-contraction coupling use the same molecular partners, STIM1 and STIM2. J Biol Chem 285, 22437–22447, doi:10.1074/jbc.M110.118984 (2010).

70 Rui, Y., Pollitt, S. L., Myers, K. R., Feng, Y. & Zheng, J. Q. Spontaneous Local Calcium Transients Regulate Oligodendrocyte Development in Culture through Store-Operated Ca(2+) Entry and Release. eNeuro 7, doi:10.1523/ENEURO.0347-19.2020 (2020).

71 Berna-Erro, A., Jardin, I., Salido, G. M. & Rosado, J. A. Role of STIM2 in cell function and physiopathology. J Physiol 595, 3111–3128, doi:10.1113/jp273889 (2017).

72 Dietz, K. C., Polanco, J. J., Pol, S. U. & Sim, F. J. Targeting human oligodendrocyte progenitors for myelin repair. Experimental neurology 283, 489–500, doi:10.1016/j.expneurol.2016.03.017 (2016).

73 Kar, P., Bakowski, D., Di Capite, J., Nelson, C. & Parekh, A. B. Different agonists recruit different stromal interaction molecule proteins to support cytoplasmic Ca2+ oscillations and gene expression. Proc Natl Acad Sci U S A 109, 6969–6974, doi:10.1073/pnas.1201204109 (2012).

74 Thiel, M., Lis, A. & Penner, R. STIM2 drives Ca2+ oscillations through store-operated Ca2+ entry caused by mild store depletion. J Physiol 591, 1433–1445, doi:10.1113/jphysiol.2012.245399 (2013).

75 Harriss, D. R., Marsh, K. A., Birmingham, A. T. & Hill, S. J. Expression of muscarinic M3-receptors coupled to inositol phospholipid hydrolysis in human detrusor cultured smooth muscle cells. J Urol 154, 1241–1245 (1995).

76 Doherty, A. J., Palmer, M. J., Henley, J. M., Collingridge, G. L. & Jane, D. E. (RS)-2-chloro-5-hydroxyphenylglycine (CHPG) activates mGlu5, but no mGlu1, receptors expressed in CHO cells and potentiates NMDA responses in the hippocampus. Neuropharmacology 36, 265–267, doi:10.1016/s0028-3908(97)00001-4 (1997).

77 Pennington, L. D. et al. 4-Methoxy-N-[2-(trifluoromethyl)biphenyl-4-ylcarbamoyl]nicotinamide: A Potent and Selective Agonist of S1P1. ACS Med Chem Lett 2, 752–757, doi:10.1021/ml2001399 (2011).

78 Vassallo, R. R., Jr., Kieber-Emmons, T., Cichowski, K. & Brass, L. F. Structure-function relationships in the activation of platelet thrombin receptors by receptor-derived peptides. The Journal of biological chemistry 267, 6081–6085 (1992).

79 Calleri, E. et al. Frontal affinity chromatography-mass spectrometry useful for characterization of new ligands for GPR17 receptor. J Med Chem 53, 3489–3501, doi:10.1021/jm901691y (2010).

80 Kawanabe, Y., Okamoto, Y., Enoki, T., Hashimoto, N. & Masaki, T. Ca(2+) channels activated by endothelin-1 in CHO cells expressing endothelin-A or endothelin-B receptors. American journal of physiology. Cell physiology 281, C1676–1685, doi:10.1152/ajpcell.2001.281.5.C1676 (2001).

81 Chhatriwala, M. et al. Induction of novel agonist selectivity for the ADP-activated P2Y1 receptor versus the ADP-activated P2Y12 and P2Y13 receptors by conformational constraint of an ADP analog. The Journal of pharmacology and experimental therapeutics 311, 1038–1043, doi:10.1124/jpet.104.068650 (2004).

82 McGuire, J. J., Saifeddine, M., Triggle, C. R., Sun, K. & Hollenberg, M. D. 2-furoyl-LIGRLO-amide: a potent and selective proteinase-activated receptor 2 agonist. The Journal of pharmacology and experimental therapeutics 309, 1124–1131, doi:10.1124/jpet.103.064584 (2004).

83 Tabarean, I. V. Functional pharmacology of H1 histamine receptors expressed in mouse preoptic/anterior hypothalamic neurons. Br J Pharmacol 170, 415–425, doi:10.1111/bph.12286 (2013).

84 Console-Bram, L., Brailoiu, E., Brailoiu, G. C., Sharir, H. & Abood, M. E. Activation of GPR18 by cannabinoid compounds: a tale of biased agonism. Br J Pharmacol 171, 3908–3917, doi:10.1111/bph.12746 (2014).

85 Hinz, S., Lacher, S. K., Seibt, B. F. & Muller, C. E. BAY60-6583 acts as a partial agonist at adenosine A2B receptors. The Journal of pharmacology and experimental therapeutics 349, 427–436, doi:10.1124/jpet.113.210849 (2014).

86 Meis, S. et al. NF546 [4,4’-(carbonylbis(imino-3,1-phenylene-carbonylimino-3,1-(4-methyl-phenylene)-car bonylimino))-bis(1,3-xylene-alpha,alpha’-diphosphonic acid) tetrasodium salt] is a non-nucleotide P2Y11 agonist and stimulates release of interleukin-8 from human monocyte-derived dendritic cells. J Pharmacol Exp Ther 332, 238–247, doi:10.1124/jpet.109.157750 (2010).

