## Supplemental Data for "Oscillatory calcium release and sustained store-operated oscillatory calcium signaling prevents differentiation of human oligodendrocyte progenitor cells"

**Extended Data:**

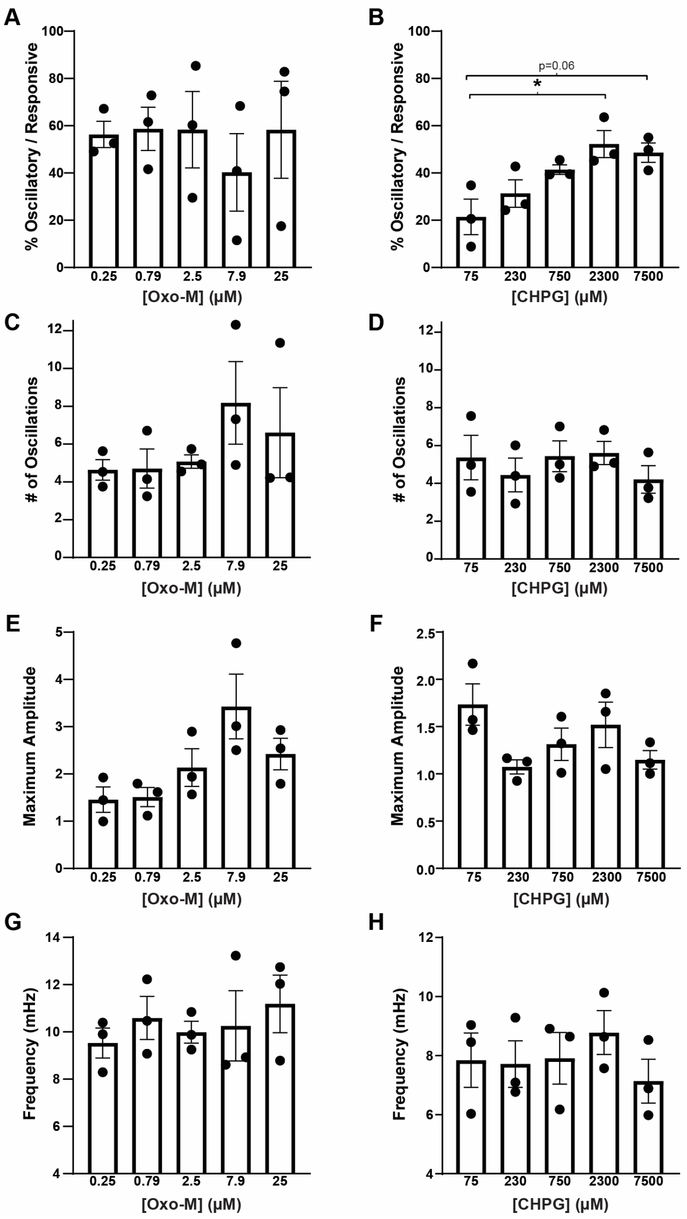


**Supplemental Figure S1 (related to Fig. 1): Comparison of oscillatory calcium signaling following $M_{1/3}R$ and $mGluR_5$ activation.** hOPCs were initially infected with GCaMP6s lentivirus prior to time-lapse microscopy in normal growth media. Assessment of oscillatory calcium signaling patterns in hOPCs *in vitro* following dose-dependent activation of $M_{1/3}R$ or $mGluR_5$ with Oxo-M or CHPG respectively. For each biological replicate preparation, all doses of either Oxo-M or CHPG were each tested on naïve cultures (1 dose/well/drug), with all doses tested on the day of experiment. Concentration specific analyses of % oscillatory responsive cells (Oxo-M: **A,** CHPG: **B**), number of calcium oscillations (Oxo-M: **C,** CHPG: **D**), maximum peak amplitude (Oxo-M: **E,** CHPG: **F**), and oscillatory response frequency (Oxo-M: **G,** CHPG: **H**). Mean ± SEM of data representative of averaged oscillatory cell responses (presenting ≥ 2 peaks) from each of 3 independent experiments (n=3 human fetal sample culture preparations). ≥ 50 cells quantified between two imaging fields per each dose/biological replicate. * $p<0.05$, RM one-way ANOVA with Holm-Sidak's post-hoc test).

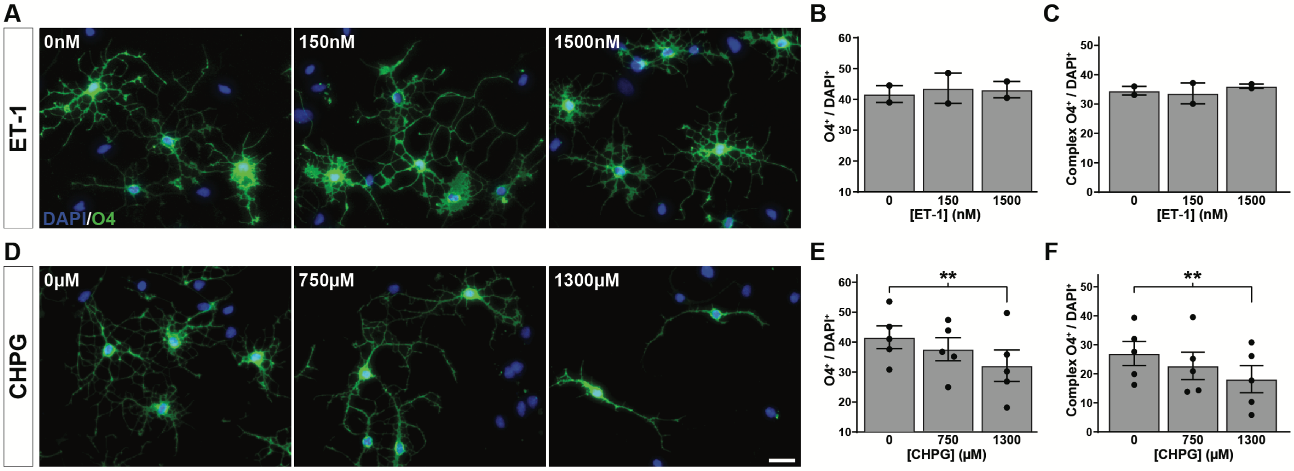


**Supplemental Figure S2 (related to Fig. 1): mGluR$_5$ activation attenuates hOPC differentiation.** The effects of various G$\alpha_q$-coupled receptor ligands on hOPC differentiation was assessed *in vitro*. Following removal of mitogens, hOPCs were treated with ET-1 (0 - 1500nM) to activate endothelin-B receptor. The effect of ET-1 on oligodendrocyte differentiation was determined by assessment of O4 immunofluorescence (**A**, green). Quantification of O4$^+$ oligodendrocytes (**B**) and morphologically complex (> 3 primary processes) O4$^+$ oligodendrocytes (**C**, mean ± SEM, n = 2 individual fetal samples). ET-1 stimulation did not influence O4$^+$ oligodendrocyte differentiation. Treatment with mGluR$_5$ agonist CHPG (0 - 1300 μM) significantly reduced O4$^+$ oligodendrocyte differentiation (**D-E**) and morphologically mature complex O4$^+$ cells (**F**, mean ± SEM (n = 5 individual fetal samples; ** $p<0.01$, RM one-way ANOVA with Holm-Sidak's post-hoc test). Scale: 25 μm.

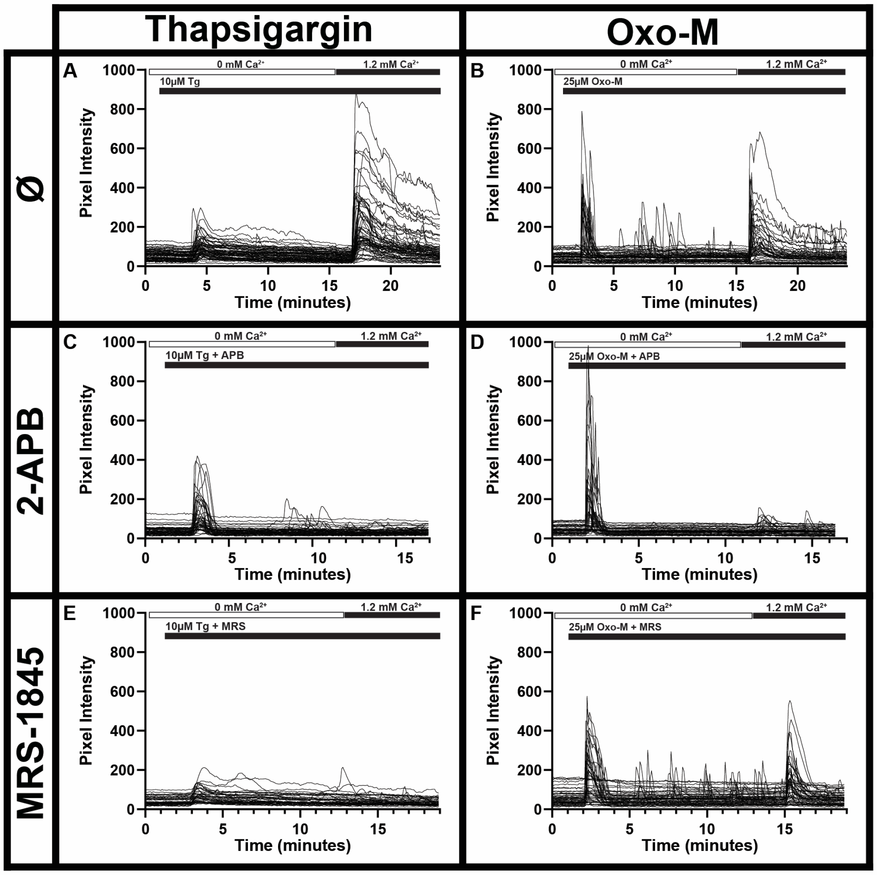


**Supplemental Figure S3 (related to Fig. 1): Per cell calcium response traces of ER depletion and SOCE following calcium re-addition**. hOPCs were initially infected with GCaMP6s lentivirus prior to time-lapse microscopy in calcium-free media. Individual per cell raw pixel intensity calcium traces corresponding to averaged traces depicted in **Fig 1G**. ER-calcium store depletion in calcium-free culture conditions and SOCE following calcium re-addition. ER-depletion induced by thapsigargin (**A**) or Oxo-M (**B**), followed by calcium re-addition and SOCE response. Blockade of SOCE following pre-incubation with SOCE antagonists 2-APB and MRS in Tg (**C, E**) or Oxo-M (**D, F**) stimulated hOPCs respectively (n ≥ 44 cells shown/quantified per condition). Timings for addition of Tg, Oxo-M and $Ca^{2+}$-containing solution are indicated by horizontal bars above each plot.

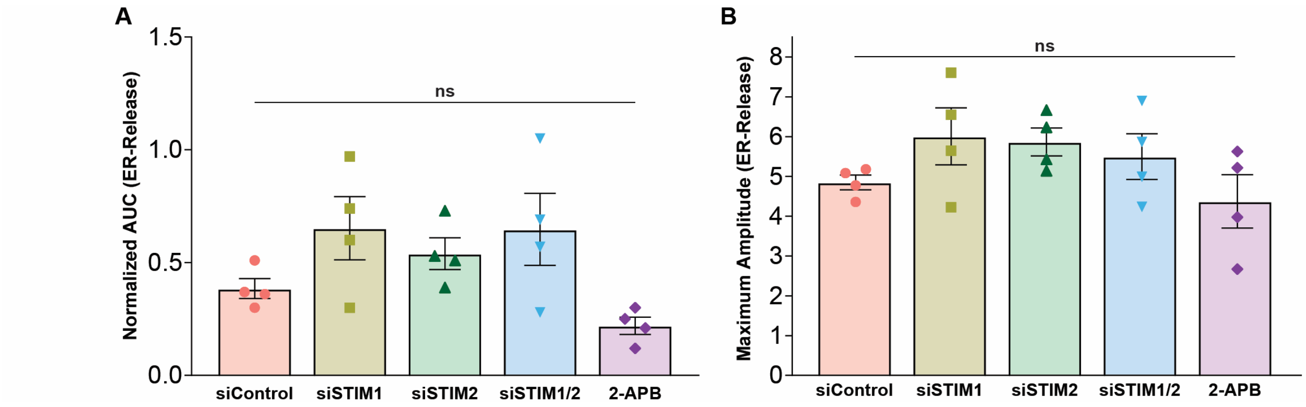


**Supplemental Figure S4 (related to Fig. 2): Effects of STIM1/2 siRNA treatment on resting hOPC ER-calcium store content.** hOPCs were initially infected with GCaMP6s lentivirus and then transfected with STIM siRNA or scrambled control siRNA, or pre-treated with 2-APB [50μM] prior to time-lapse microscopy in calcium free media. Oxo-M [25μM] was used to deplete ER-Stores after a one minute baseline, and responses measured. **A,** the ER-calcium store content was measured by quantification of the area under the curve (AUC) of the initial peak calcium response for an additional 10m post Oxo-M addition normalized to Influx of control cells within matched biological replicates. following treatment with Oxo-M [25μM] and measured for 10 minutes following Oxo-M treatment. **B.** quantification of maximum peak amplitude of ER-depletion following Oxo-M addition. Cells were analyzed across two imaging fields in a single well per each condition and all responses were averaged per condition for each biological replicate. Data are presented as mean ± SEM of averaged cell responses per each of three independent experiments (n=3 independent human fetal sample preparations), with >140 total cells quantified per each condition. RM one-way ANOVA with Holm-Sidak's post-hoc test.

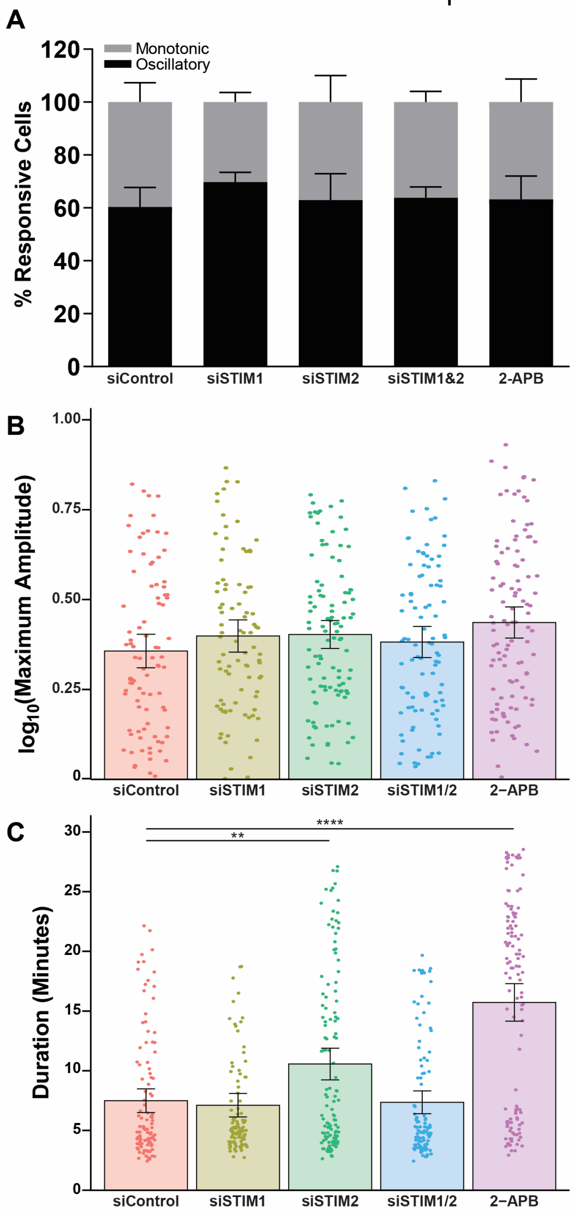


**Supplemental Figure S5 (related to Fig. 2): Effect of STIM1/2 siRNA treatment on muscarinic induced oscillatory calcium responses in hOPCs.** hOPCs were initially infected with GCaMP6s lentivirus and then transfected with STIM siRNA or scrambled control siRNA, or pre-treated with 2-APB [50μM] prior to time-lapse microscopy in normal growth media. **A**, STIM1/2 KD does not influence the total percentage of oscillatory responsive hOPCs following muscarinic stimulation (RM one-way ANOVA, $p>0.45$). **B**, STIM1/2 KD does not affect the maximum peak amplitude ($p=0.4$) of Oxo-M induced calcium responses in hOPCs. Data represent $Log_{10}$transformed mean ± SEM of averaged cell responses per each of four independent experiments ($n=4$ independent human fetal sample preparations). **C**, STIM2 KD and 2-APB treatment increased the oscillatory response duration of muscarinic induced calcium signaling in hOPCs (Mean ± SEM, $n=4$) . Cells were analyzed across two imaging fields in a single well per each condition and all responses were averaged per condition for each biological replicate, with >100 total cells quantified per each condition. ** $p<0.01$, ****$p<0.0001$, linear model with Tukey's HSD posttest.

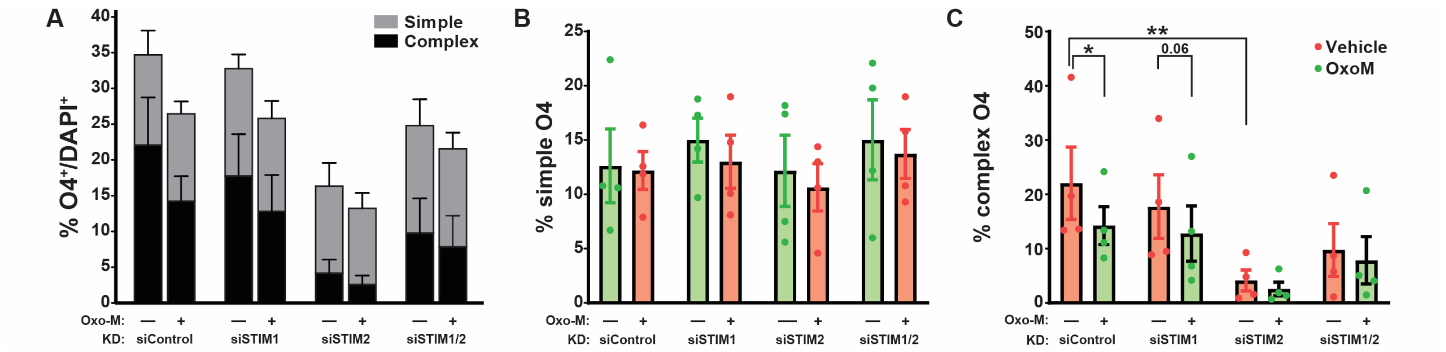


**Supplemental Figure S6 (related to Fig. 3): Effect of STIM1/2 siRNA treatment of human oligodendrocyte morphological maturation.** Morphological maturation of $O4^+$ oligodendrocytes was assessed by determining the proportion of complex branching and simple process bearing cells. $O4^+$ oligodendrocytes were characterized as complex if they had at least 3 highly branched processes. **A**, quantification of both simple and complex $O4^+$ oligodendrocytes. Mean ± SEM shown (n=4 fetal samples). **B**, analysis of simple $O4^+$ oligodendrocytes. Three-way ANOVA indicates that STIM1 or STIM2 do not influence the percentage of simple $O4^+$ oligodendrocytes (main effects, $p>0.3$). **C**, analysis of complex $O4^+$ oligodendrocytes. Three-way ANOVA using Oxo-M, STIM1 and STIM2 as factors revealed a significant effect of STIM2 siRNA on complex $O4^+$ cells ($F$ (1, 3) = 28.64, $p=0.02$). * $p<0.05$ and ** $p<0.01$ pairwise Holm-Sidak's post-hoc test as indicated.
